# The emergence of the language system in the toddler brain

**DOI:** 10.64898/2026.02.26.707550

**Authors:** Halie A. Olson, Emily M. Chen, Camille J. Osumah, Haoyu Du, Evelina Fedorenko, Rebecca Saxe

**Affiliations:** McGovern Institute for Brain Research, Massachusetts Institute of Technology; Department of Brain and Cognitive Sciences, Massachusetts Institute of Technology; Department of Psychology, Stanford University

## Abstract

Toddlerhood is a time when children’s language competence undergoes a dramatic transformation, making it a key period to examine the emergence of the brain’s language system. Functional magnetic resonance imaging data from 29 awake toddlers (19-36 months) scanned on a child-friendly language task reveal that, already at this young age, areas in the left frontal and temporal cortex respond more to comprehensible language (i.e., videos of puppets speaking in English) than the control condition (matched videos with incomprehensible audio). Although the magnitude of response to language is substantially weaker than in adults, the topography is already adult-like, challenging claims that the frontal component or left-hemispheric dominance do not emerge until later in life.

---

Language is a remarkable capacity of the human brain, surpassing other animal communication systems in complexity and generative power. Language allows humans to share memories of the past and aspirations for the future, talk about imaginary and implausible things, and teach each other new knowledge and skills. In the adult brain, this capacity draws on a specialized system of temporal and frontal areas in the left hemisphere ^[1–6]^. These areas respond strongly during language comprehension and production across modalities (i.e., spoken, signed, written), but are not engaged in other types of cognition (for review, see ^[2]^). When and how does this specialized system emerge in ontogeny?

From a behavioral standpoint, language development follows a protracted trajectory over the first years of life. Young infants learn to discriminate and produce the sounds of their native language, but they only begin to show hints of mapping common words to meaning around 6 months of age ^[7]^. From there, infants slowly learn additional word-meaning associations and can produce a handful of spoken words by their first birthday. Then a remarkable leap occurs, and toddlers begin rapidly acquiring words and larger constructions, such that by two years later, by three years of age, they understand and produce hundreds of words and word combinations ^[8–10]^.

From a neural standpoint, the trajectory of language network development in the brain is less clear. Multiple studies have examined brain responses to speech sounds in infants and newborns (e.g., ^[11–19]^), but until infants begin linking word forms to meanings, it is impossible to examine ‘language’ per se (in contrast with speech, which engages a separate system in the mature human brain – a system that responds to speech sounds regardless of whether they convey meaning ^[20]^). To understand the emergence of the *language* system in the brain, one needs to examine children at an age when they becoming capable of linguistic communication, which happens during toddlerhood. One possibility is that toddlers will not yet show robust, measurable language responses when controlling for non-linguistic features that typically co-occur with spoken language (e.g., recognizable agents who are speaking, acoustic features of speech, communicative signals, social contingency), particularly since toddlers only know tens to hundreds of words ^[8]^ and thus much of their language input is incomprehensible to them. An alternative possibility is that, despite their limited comprehension, toddlers will already show the beginnings of selective responses to language, in approximately the same cortical areas that support language processing in adults.

If a language system exists in some form in toddlers, how might it differ from the mature language network? Two broad ideas have been put forward about how the topography of brain responses changes during early language development. The first hypothesis is that language initially recruits bilateral cortical areas and gradually becomes left-lateralized (e.g., ^[21]^). This hypothesis was inspired by evidence that damage to the left hemisphere early in life can have little or no impact on language skills (^[22–25]^; *cf*. ^[26,27]^), in sharp contrast to similar damage in adulthood where aphasia typically results (e.g., ^[28,29]^). To explain this observation, researchers hypothesized that both hemispheres support language processing early in life, with a gradual reduction of the right hemisphere activity with age ^[21,30]^.

The second broad hypothesis is that cortical development roughly follows a posterior-to-anterior developmental hierarchy (e.g., ^[31]^). This hypothesis is grounded in the generally protracted maturation of the frontal lobes, for example as measured by gray matter volume, myelination, and synaptic density (e.g., ^[32–34]^). Applied to language, this hypothesis predicts that language processing should initially engage only the temporal areas and then gradually incorporate frontal regions later in development ^[35]^.

Each of these hypotheses has received some empirical support (e.g., initially more bilateral language processing: ^[30,36,37]^; initially temporal-lobe-dominant language processing: ^[37– 40]^), but evidence has also accumulated against both of these ideas, with children showing a strong left-hemispheric bias and robust responses to language in both temporal and frontal lobes ^[30,40–44]^. However, the youngest children in prior neuroimaging studies of language are 3-5 years old, meaning that the “heavy lifting” of acquiring a language has already happened.

To test these two broad hypotheses about the early topography of responses to language in human development, we examined responses to language in the brains of awake toddlers using fMRI. Compared to fNIRS and EEG, two other noninvasive neuroimaging tools more often used with infants and toddlers, fMRI offers whole-brain coverage and high spatial resolution ^[45]^. On the other hand, fMRI in awake toddlers presents a truly formidable methodological challenge ^[46,47]^. How do you get a 2-year-old, who may be undergoing a period of high anxiety and novelty avoidance ^[48]^, who has only recently learned to locomote ^[49]^, and whose executive mechanisms are highly immature ^[50]^, to get into a scary and loud machine, remain still, and pay attention for a sustained period of time? We overcame these challenges by (i) implementing a toddler-friendly preparation and scanning protocol (**Fig. 1A, B**), and (ii) using as stimuli clips from *Sesame Street* – a television show with decades of programming explicitly designed for the target age group and which has already proven effective in developmental neuroimaging research ^[51]^. The language stimuli are video clips in which a character speaks directly to the child (“monologue”, akin to a caretaker talking to the child) and video clips in which two characters speak to each other (“dialogue”, akin to observing others’ linguistic exchanges). Importantly, to isolate neural responses to language from responses to incomprehensible speech while sustaining toddlers’ attention and engagement in the task, we created a control condition by reversing the audio track of characters’ speech and overlaying it on the video (**Fig. 1D**; see **Fig. S1** for behavioral evidence that the forward and backward speech videos sustain toddlers’ attention to similar degrees).

**Fig. 1.**
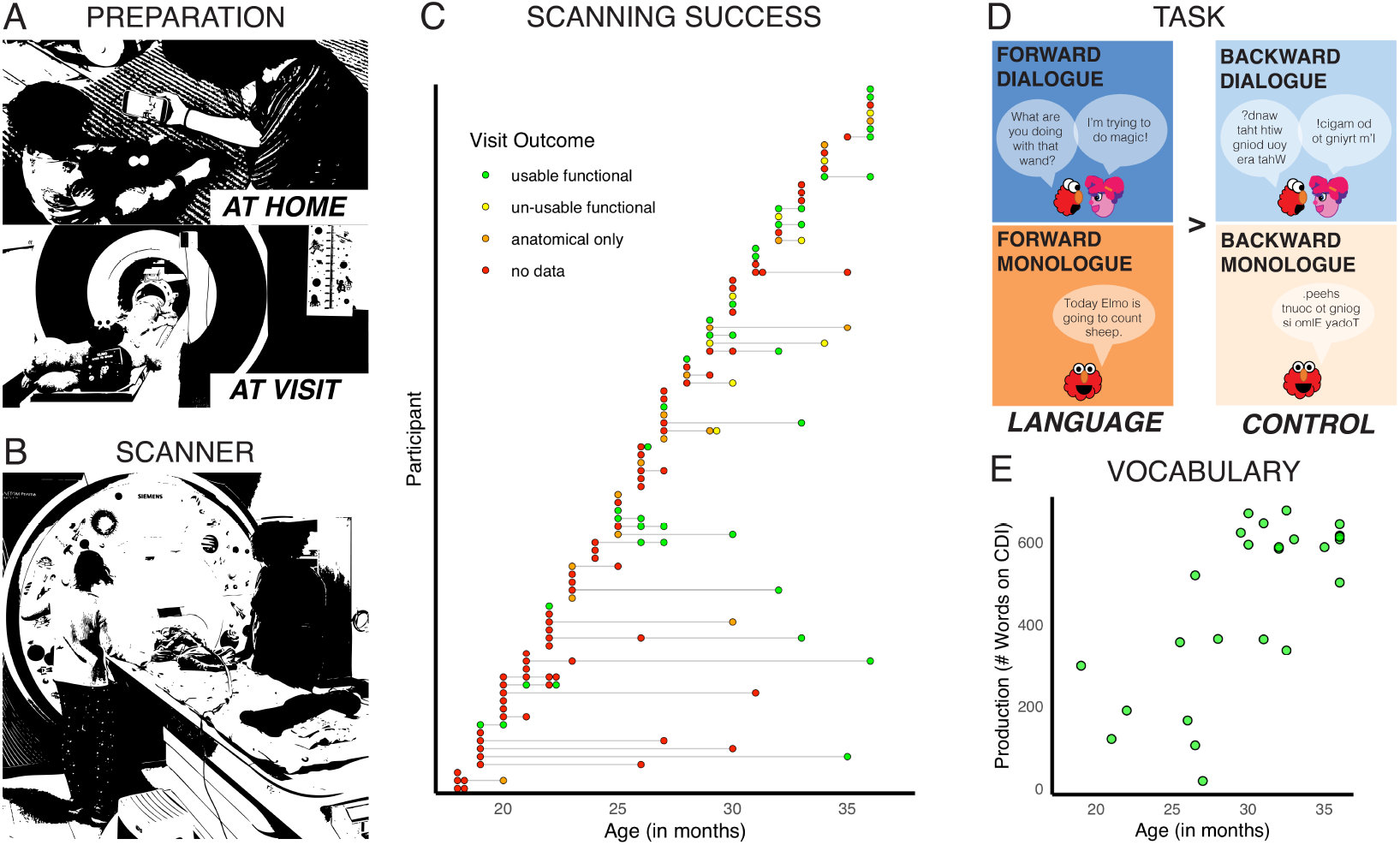
Methods overview. **(A)** Top: A parent prepares her child for the scan by watching our video at home (link: https://youtu.be/P2yGWnp7s). Bottom: Toddlers acclimate to the MRI environment in the mock scanner. **(B)** A 21-month-old toddler gets ready to go in the MRI scanner. **(C)** Age in months and visit outcomes for all participants. Each participant (total N=89) is represented by a row and each visit by a dot, with gray lines connecting multiple visits for the same participant (2-4 visits for 40% of participants). Green dots indicate that usable functional MRI data were collected and included in the final sample; yellow dots indicate that some functional MRI data were collected but not included in the final sample due to motion or other data quality issues; orange dots indicate that some anatomical, but no functional, data were collected; and red dots indicate that no MRI data were collected at the visit. **(D)** For the fMRI task, participants watched 20 seconds of videos from *Sesame Street* with one (monologue) or two (dialogue) characters at a time, with the audio either played normally (forward) or reversed by character (backward), with the main language contrast defined as Language (the two forward conditions) > Control (the two backward conditions). **(E)** Raw scores for vocabulary production (number of words endorsed by a caregiver on the MacArthur Bates Communicative Development Inventory (CDI; ^[52]^) as the child “knows and says”) for included participants are distributed across the sample and positively correlated with age (N=25 shown; 4 participants were missing CDI data and are not included in this plot).

This experimental design provides a few key advantages. First, unlike many studies that use auditory-only stimuli (e.g., forward vs. backward speech), the audiovisual modality engages toddlers’ attention and ensures that the control conditions are also communicative. The reversed speech is carefully overlaid on the video to correspond to the correct puppet, creating the impression of the characters “speaking.” Second, including monologue and dialogue conditions allows us to test whether language responses generalize across two contexts in which toddlers encounter language. Complementarily, we test – using the same stimuli – whether responses to language can be dissociated from cortical responses to the social and perceptual content of the videos, by comparing dialogue to monologue.

The data collection procedures, stopping criteria, and an analysis plan for this study were preregistered. Usable fMRI data were collected from 29 toddlers (ages 19-36 months, mean(SD)=30.07(4.77) months, 19 female). An additional 60 toddlers participated in the study but failed to provide usable data (39 with no MRI data, 13 with only anatomical MRI data, and 8 with functional MRI data that were not usable due to motion or data quality; **Fig. 1C**). Eligible toddlers had exposure to English (some also had variable levels of exposure to other languages), no known hearing or neurodevelopmental difficulties, and no MRI contraindications. Parents filled out the English (American) Words & Sentences form of the MacArthur Bates Communicative Development Inventory (CDI; ^[52]^) as a measure of productive vocabulary. As expected, vocabulary was correlated with age (*r = 0*.71; Pearson’s test for *r*>0: t = 4.80, p < 0.001; **Fig. 1E**). Twenty adult participants also completed the fMRI task and are included as a comparison group ^[53]^.

## DO TODDLERS HAVE A LANGUAGE NETWORK?

We first asked: do toddlers show selective responses to language in cortex, as early as language competence emerges, in approximately the cortical areas that adults do? If so, does this language response generalize across contexts in which toddlers encounter language: when it is directed at them, and when they are observing others talking to each other?

For the critical tests of our hypotheses, we individually defined functional regions of interest (fROIs) in canonical left hemisphere cortical language regions. Defining fROIs within individuals accounts for the well-established inter-individual variability in the locations of functional areas and helps isolate target areas from nearby functionally distinct areas ^[54–57]^. In 17 toddlers with multiple runs of the fMRI task, we iteratively defined functional regions of interest (fROIs) for language ^[54]^ by selecting the top 100 voxels for the Language>Control contrast per region using a leave-one-run-out approach (note that the results also hold when defining fROIs as the top 10% of voxels for each region; see **Table S9**). Language regions of interest included five search spaces derived from adults completing a language localizer task ^[54]^ in left orbital inferior frontal gyrus (IFGorb), inferior frontal gyrus (IFG), middle frontal gyrus (MFG), anterior temporal (AntTemp), and posterior temporal (PostTemp) cortex. We then extracted the response magnitude for each condition in the held-out run from each language fROI and averaged across leave-one-run-out iterations (**Fig. 2A**).

**Fig. 2.**
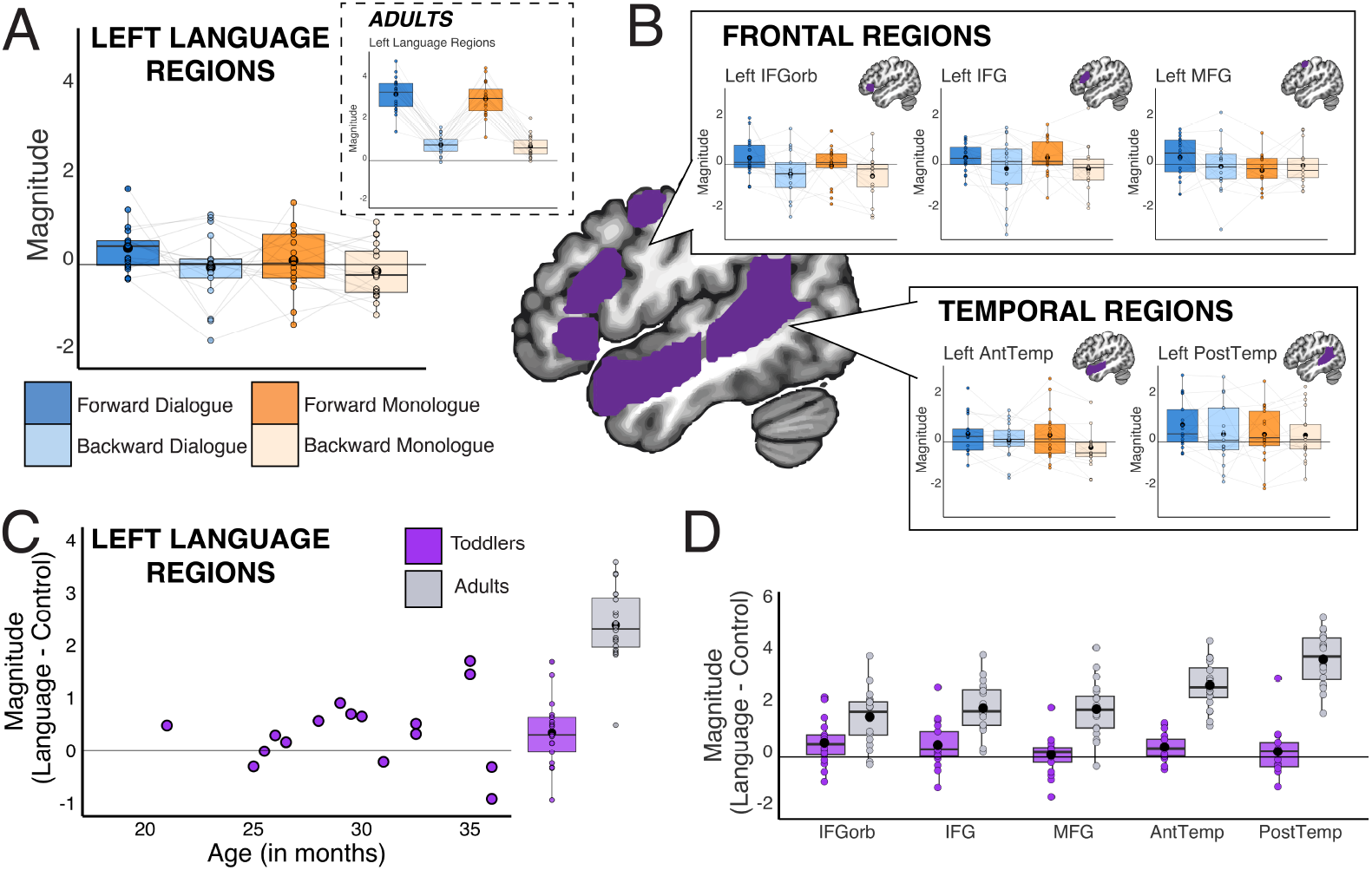
Responses to language in toddler brains. For participants with multiple usable functional runs, fROIs for the top 100 voxels were iteratively identified using the Language>Control contrast in each language parcel using a leave-one-run-out approach. **(A)** Boxplots show response magnitude for held-out runs (independent data) for each condition, in toddlers (N=17) and adults (N=20; *inset*), averaged across all five language regions. Note that the low-level baseline for adults was a silent fixation cross ^[53]^, whereas toddlers viewed audiovisual music clips ^[58]^ during baseline periods in order to sustain attention; thus, differences in response relative to baseline are not necessarily age-related. Dots show individual participants’ responses; light gray lines connect the same participant; large black dots show means. **(B)** Language search spaces are plotted on the brain in purple; boxplots show toddlers’ responses per condition for each region. **(C)** The difference in magnitude between language and the control conditions, averaged across language regions, was not associated with age in toddlers. **(D)** For each language region, we also plot the difference in magnitude between language and control conditions in each individually-defined region for toddlers (purple) and adults (gray).

As our preregistered test of whether canonical left-hemisphere language regions are selective for language in toddlers, we used a linear mixed effects model to examine the effects of language comprehensibility (forward vs. backward speech), social context (dialogue vs. monologue), and cortical lobe (temporal lobe vs. frontal lobe) on univariate responses in these individually-defined language regions, accounting for age. Overall, the canonical left-hemisphere language regions showed higher responses to language than the control conditions in toddlers (Forward>Backward: Est = 0.175, S.E. = 0.050, t = 3.492, p = 0.0005), as expected for areas that are engaged in language processing. These regions did not show sensitivity to the social context (Dialogue>Monologue: Est = 0.094, S.E. = 0.050, t = 1.869, p = 0.062); thus, the language response generalized across two contexts in which toddlers encounter language.

Temporal regions showed larger responses to all stimuli than frontal regions (Temporal>Frontal: Est = 0.162, S.E. = 0.060, t = 2.702, p = 0.047), but there was no interaction between cortical lobe and either functional contrast. Finally, age was not significant in the model (Est = 0.005, S.E. = 0.024, t = 0.200; p = 0.844). The magnitude of response to the language versus control conditions, averaged across the individually-defined left-hemisphere language regions, was also not correlated with age in toddlers (**Fig. 2C**; *r = 0*.10; Pearson’s test for *r* ≠ 0: t = 0.39, p = 0.70). This analysis confirmed that the canonical left-hemisphere language network already shows sensitivity to language in toddlerhood.

We also examined responses within individual language regions of interest (**Fig. 2B**). A significantly stronger response to language than the control condition was present in one frontal region, left IFGorb (Forward>Backward: Est = 0.288, S.E. = 0.104, t = 2.774, p = 0.008, uncorrected), and one temporal region, AntTemp (Forward>Backward: Est = 0.204, S.E. = 0.064, t = 3.171, p = 0.003, uncorrected). A marginally stronger response to language was found in another frontal region, IFG (Forward>Backward: Est = 0.242, S.E. = 0.110, t = 2.200, p = 0.032, uncorrected). The language contrast was not significant in MFG or PostTemp. Just as in the model across the left hemisphere language network, Dialogue>Monologue, interactions, and age were not significant in individual regions’ models (see **Table S4** for full model results for individual regions).

## HOW DOES THE LANGUAGE SYSTEM DIFFER BETWEEN TODDLERS AND ADULTS?

Toddlers have an emerging language system in the left hemisphere of cortex, but it is not yet fully adult-like. In particular, one way that toddlers’ language responses differ from adults’ is in magnitude. Both in the language network and for each individual language region, toddlers showed weaker language responses than adults (**Fig. 2C, D**). Do toddlers’ language responses also differ in their topography? We examined key topographic features of the mature language system – left lateralization and the involvement of both frontal and temporal regions – which had been hypothesized to differ in development ^[21,30,35]^.

To examine lateralization of the language response, we considered both response magnitude and extent of activation across hemispheres. In older children and adults, both the magnitude of the language response and the extent of language-evoked responses are greater in the left hemisphere than the right hemisphere ^[42]^. Having already found that the left hemisphere language regions showed a significant response to language, we first tested whether right hemisphere homotopes of the language network are language-selective in toddlers. Right hemisphere homotopes of the canonical left-hemisphere language regions (using mirror parcels of the five canonical regions in the right hemisphere) did not show an effect of language comprehension (Forward>Backward: Est = 0.065, S.E. = 0.050, t = 1.296, p = 0.196) or social context (Dialogue>Monologue: Est = 0.049, S.E. = 0.050, t = 0.980, p = 0.328); see **Supplementary Text** for additional model terms and individual regions. Further, the size of the Language>Control effect is more than twice larger in the left hemisphere (Est = 0.175) than in the right hemisphere (Est = 0.065).

To statistically test for the presence of a hemispheric bias, we turned to the commonly used laterality index (LI), which is based on the extent of activation. In this analysis, we used all toddler participants with at least one usable run (N=29), increasing the power of this analysis.

We calculated LI for language by comparing the proportion of voxels significantly activated by language (Language>Control contrast) in the left hemisphere canonical language parcels and the mirrored right hemisphere homotope parcels in each participant (**Fig. 3A**; threshold: Z>1.64, cluster threshold k>=10; see **Table S6** for similar results at different thresholds). Overall, LI was significantly greater than 0, confirming that language responses were left-lateralized in toddlers (mean(SD)= 0.25(0.61); one-sample t-test for LI>0: t = 2.28, p = 0.015). Language responses were also left-lateralized in the comparison sample of adults, as expected (**Fig. 3A**; mean(SD)= 0.23(0.16); one-sample t-test for LI>0: t = 6.33, p <0.001). Lateralization did not differ between toddlers and adults (Welch two sample t-test: t = −0.229, p = 0.820). Moreover, LI was not correlated with age in toddlers (*r = 0*.08; Pearson’s test for *r*>0: t = 0.431, p = 0.335). A caveat, however, was that LI values were quite noisy in individual toddler participants (*c*.*f*. adults, ^[59]^), with no evidence of consistent LI scores within individuals across runs (see **Supplementary Text**). Together, both the fROI and LI analyses suggest that the language response is already left-lateralized in toddlerhood.

**Fig. 3.**
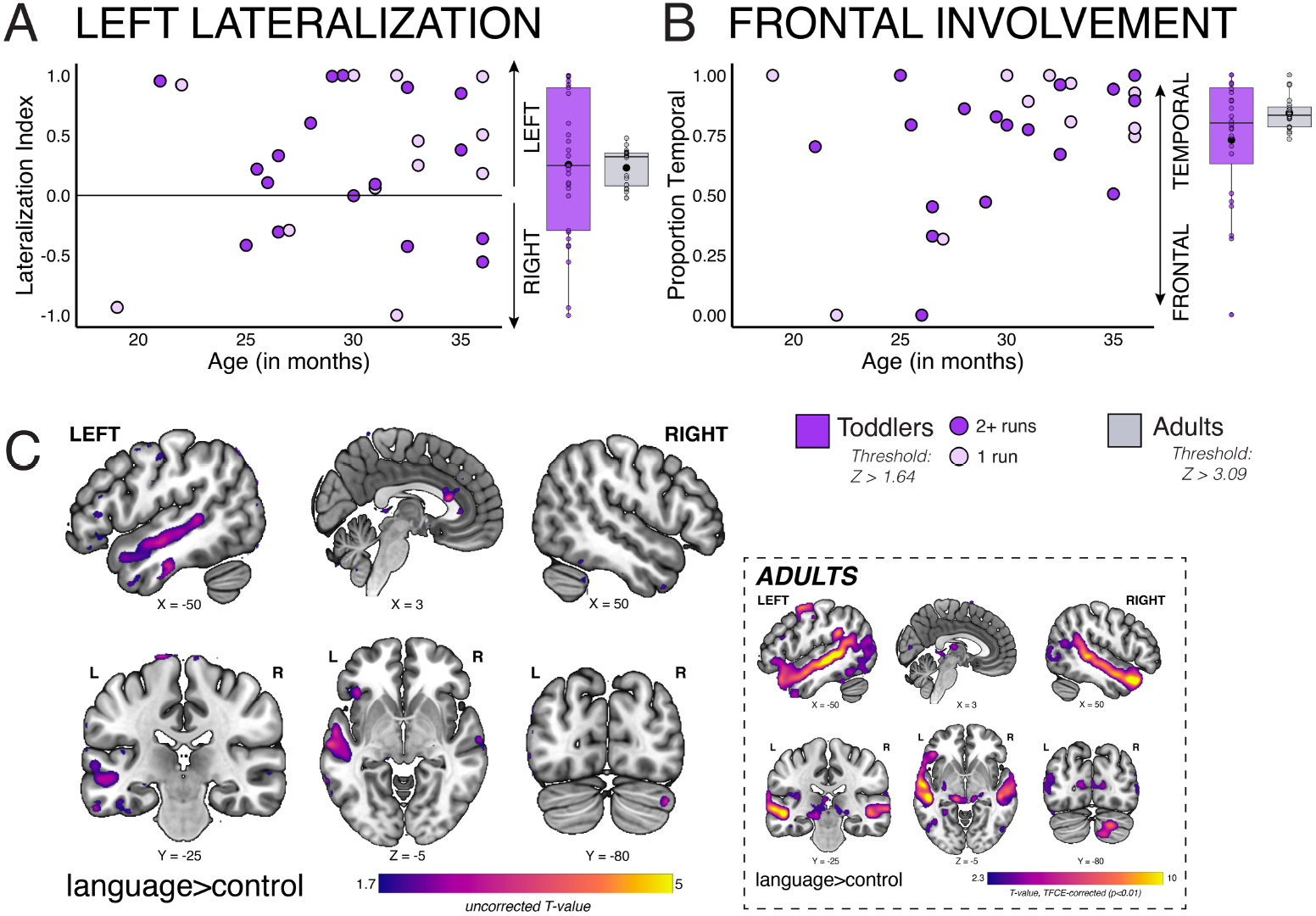
Language topography is similar between toddlers and adults. **(A)** For all 29 toddlers, lateralization index was calculated for language, using threshold Z>1.64, cluster threshold 10 voxels to identify suprathreshold voxels for the language contrast (Language>Control) in the left hemisphere language parcels and the mirror of these parcels in the right hemisphere. Lighter purple dots indicate toddlers with only one run of usable data. In adults (N=20), we applied the same approach to calculate lateralization for language, using a threshold of Z>3.09, cluster threshold 10 voxels (gray boxplot). **(B)** In the left hemisphere language parcels, the proportion of suprathreshold language voxels in the temporal regions was also calculated using the same threshold as lateralization index (NA=1 with no suprathreshold voxels). **(C)** A group random effects analysis was conducted in the full sample of toddlers for the language contrast, visualized at an uncorrected threshold T>1.70. A group random effects analysis was also conducted in adults, visualized at TFCE-corrected threshold of p<0.01 (*inset*).

A second topographic feature of the mature language system is that it includes both temporal and frontal regions. To examine the degree of frontal involvement in language processing in toddlers, we conducted an exploratory analysis inspired by the LI measure of lateralization. In particular, we examined the proportion of language voxels in the temporal language areas relative to all language-responsive voxels in the left hemisphere language regions (proportion temporal; **Fig. 3B**). For toddlers, mean(SD)= 0.729(0.28) (one-sample t-test for proportion temporal<1: t = −4.967, p<0.001; one participant was excluded due to no LH suprathreshold voxels). The proportion of temporal voxels was similarly high in adults: mean(SD)= 0.841(0.071) (one-sample t-test for proportion temporal<1: t = −10, p <0.001), with no significant difference between the age groups (Welch two sample t-test: t = 1.974, p = 0.057). Moreover, the proportion of temporal voxels was not correlated with age in toddlers (*r =* 0.376; Pearson’s test for *r*<0: t = 2.069, p = 0.976). These results suggest that language system of toddlers already includes a frontal component, and the temporal-to-frontal distribution is similar to that of adults.

Finally, as a complementary test of the spatial distribution of language-sensitive responses and to test whether the toddler group may show activity in some additional brain areas, we conducted an unbiased whole brain group analysis across all toddlers (N=29) using the Language>Control contrast ([forward dialogue + forward monologue]>[backward dialogue + backward monologue]). The whole-brain random effects analysis at the preregistered corrected threshold (TFCE corrected, p<0.01) did not identify any significant clusters. Whole-brain group analyses have low sensitivity, and this lack of significant group differences may reflect inter-individual variability in the precise locations of brain areas ^[57,60]^. At a more lenient threshold (T>1.70, uncorrected; **Fig. 3C**), several clusters emerged that responded more to language than the control conditions, resembling the canonical language network, including left posterior and anterior temporal cortex and right Crus I of the cerebellum (see **Table S2** for list of significant clusters at this exploratory uncorrected threshold). The adult comparison group showed responses in all regions of the canonical language network (**Fig. 3C**). These results show that responses to language in toddlers are restricted to brain areas that respond to language in adults.

## ARE LANGUAGE AND SOCIAL PROCESSING DISSOCIATED IN THE TODDLER BRAIN?

Our experimental design allowed us to explore one other distinction that is robust in adults: between regions that respond to language and those that respond to the visual and social content of the videos. Given that language regions already show a left-hemisphere bias, we tested whether social processing likewise shows a right-hemisphere bias in toddlers. In an exploratory analysis, we calculated LI for the social processing (Dialogue>Monologue) contrast within the same parcels given that socially-responsive areas tend to exhibit similar fronto-temporal topography ^[61]^. In adults, the Dialogue>Monologue response was right-lateralized in these regions (mean(SD)=-0.31(0.44); one-sample t-test for LI<0: t = −3.23, p = 0.002). In toddlers, the Dialogue>Monologue response was trending toward right-lateralization (**Fig. 4A**; mean(SD)=-0.19(0.60); one-sample t-test for LI<0: t = −1.63, p = 0.058) and became more right-lateralized with age (*r =*-0.41; Pearson’s test for *r*≠0: t = −2.26, p = 0.03).

**Fig. 4.**
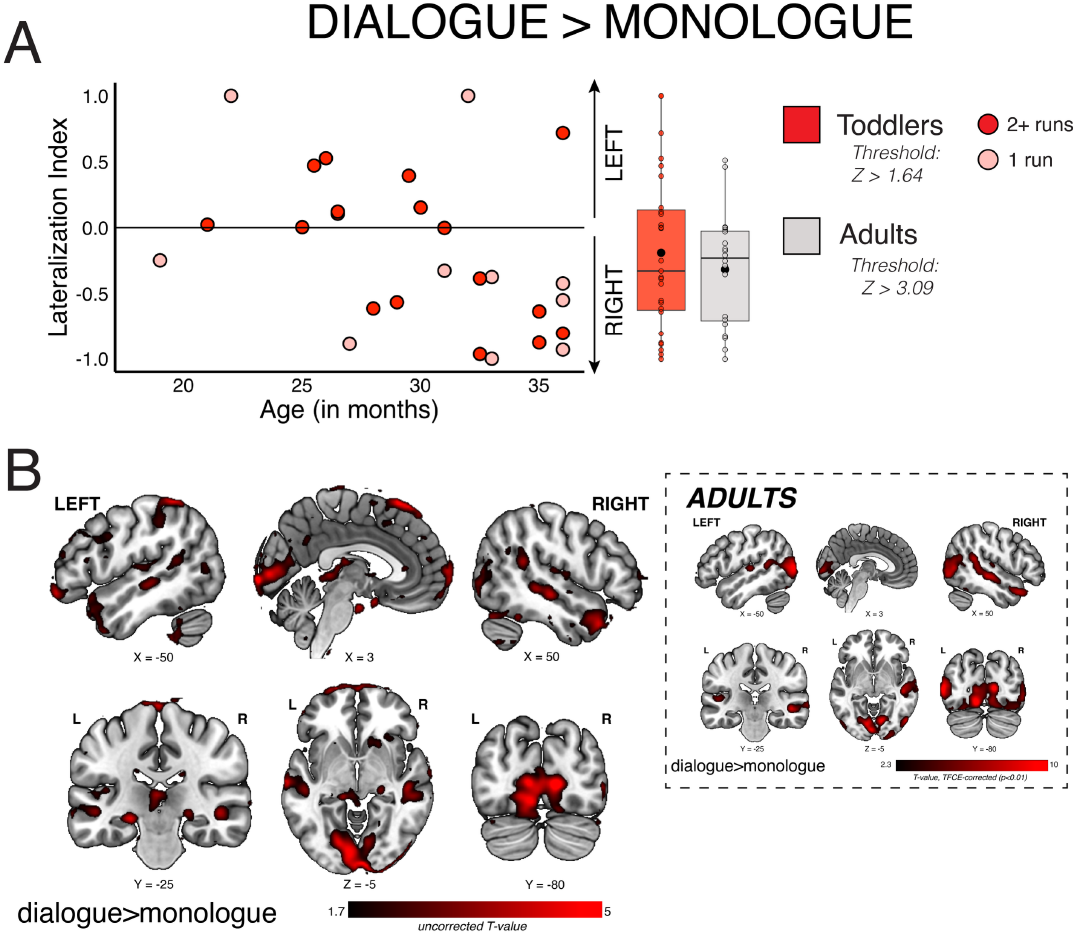
Responses to social processing. **(A)** For all 29 toddlers, lateralization index was calculated for a social contrast, using threshold Z>1.64, cluster threshold 10 voxels to identify suprathreshold voxels for Dialogue>Monologue in the left hemisphere language parcels and the mirror of these parcels in the right hemisphere (NA=2 with no suprathreshold voxels). Lighter dots indicate toddlers with only one run of usable data. In adults (N=20) we applied the same approach, using a threshold of Z>3.09, cluster threshold 10 voxels (gray boxplot). **(B)** A group random effects analysis was conducted in the full sample of toddlers (N=29) for the Dialogue>Monologue contrast, visualized at an uncorrected threshold T>1.70. A group random effects analysis was also conducted in adults, visualized at TFCE-corrected threshold of p<0.01 (*inset*).

We also conducted a whole-brain random effects analysis for the Dialogue>Monologue contrast, which revealed similar regions in both toddlers and adults (**Fig. 4B**), including, as expected, superior temporal cortex, as well as bilateral occipital cortex, presumably due to differential visual demands, which were not matched between these conditions (see **Table S3** for significant clusters).

## DISCUSSION

To understand the origins of the brain’s language system, we need to study it when linguistic behavior emerges in the first years of life ^[62]^. Here we found that the basic functional architecture for the language system is already present during toddlerhood, a critical developmental stage that encompasses the transition from understanding a few isolated words to understanding and expressing complex meanings through language ^[8–10]^. Toddlers in our study averaged just a few hundred spoken words, a tiny fraction of the 20-50 thousand words they will know by adulthood ^[63]^. Yet like older children and adults ^[1–5]^, 19-36-month-old toddlers recruited canonical frontal and temporal cortical regions when comprehending language. This response was lateralized to the left hemisphere, and as in adults ^[53]^, generalized across manipulations of social context (i.e., dialogue vs. monologue). The topography of this language-evoked response differed from toddlers’ responses to viewing social interactions. These results help fill a key gap in our understanding of the ontogenetic emergence of the language system, demonstrating continuity in the spatial and functional characteristics of cortical regions supporting language comprehension from toddlerhood onwards.

### Left lateralization of toddlers’ language responses

Our results provide evidence against two notable hypotheses about the neural underpinnings of language development. The first hypothesis is that language responses emerge bilaterally and become more left-lateralized in cortex across development ^[21]^. Prior evidence for this hypothesis has not only been mixed, but also limited to either older children who had already acquired fluency in language (e.g., ^[30,36,42]^) or infants who did not yet comprehend language (e.g., ^[11,16,17,19,64]^). Here, we addressed a critical gap in the developmental timeline by examining toddlers, whose language comprehension is in a critical early stage, finding that language-evoked activation was already left-lateralized and not associated with age.

What does it mean for toddlers to already show a left-hemisphere bias for language?

Laterality is a ubiquitous feature of the vertebrate brain from genetic expression to cortical areas ^[65–67]^. Many brain asymmetries develop before birth ^[68]^, and prematurity in humans is associated with atypical lateralization ^[69]^. Thus, it is perhaps unsurprising that laterality for language comprehension is present as early as we can measure it. An intriguing possibility is that early lateralization for language may be beneficial for language learning. One of the biggest puzzles in language development is how young children can so quickly and efficiently learn language, which has been underscored by the massive amounts of training data needed for large language models to achieve the same feat ^[70]^. Lateralization has been shown to confer a processing efficiency benefit across species and systems ^[71–73]^, so it is possible that the left-lateralized organization of the emerging language system is more efficient than a bilateral system. That said, why the left hemisphere specifically, rather than the right one, is language-dominant in most individuals remains an open question.

It is also worth considering the other side of this lateralization puzzle. The corollary of language in the left hemisphere may be a bias toward social functions in the right hemisphere ^[61]^. In the same regions that showed left lateralization for language, we found that the response to dialogue vs. monologue was right lateralized in adults and showed a similar bias in toddlers. Furthermore, responses for the dialogue>monologue contrast became more right-lateralized across age in toddlers (however, we note that the dialogue and monologue conditions differ along multiple dimensions and thus the contrast does not cleanly isolate social processing).

Together, this evidence points to both a clear pattern of lateralization as language is emerging in toddlerhood, as well as an early dissociation between language and social processing in the developing brain.

### Frontal involvement

The second hypothesis we examined is that language specialization in the brain follows a temporal-to-frontal developmental trajectory ^[35]^. At least some version of this hypothesis might predict that toddlers would only show language-specific responses in temporal regions. Instead, frontal areas (left IFGorb and IFG) were also more engaged for comprehensible language than the control conditions in toddlers. These findings are consistent with evidence that the prefrontal cortex is already engaged in cognitive processing in infancy ^[74]^, as well as evidence that frontal areas are involved in early speech processing (e.g., ^[11,14,75]^).

The precise role of these emerging frontal language-sensitive areas in toddlers is yet to be determined. Do they support similar computations as the temporal language regions during the early stages of language acquisition? By adulthood, the frontal and temporal language regions form a cohesive whole ^[2,54,76]^. This may or may not be the case for the emerging language network in toddlers. Recent evidence from stereotactic EEG, for instance, revealed that word-level representations could be detected in 2-5-year-old participants’ superior temporal gyrus but not lateral/inferior frontal cortex ^[77]^. Thus, it is possible that frontal regions are supporting language comprehension in toddlers in a different way than temporal regions early in language development, but more data will be needed to understand these potential differences.

### Lower response magnitude in toddlers

The difference between the comprehensible language conditions and the control conditions was much smaller in toddlers compared to adults. Multiple factors likely contributed to the smaller hemodynamic contrast in toddlers. First, we know that hemodynamic responses change across early development, even when neural responses are constant, and BOLD responses are often lower in infants and toddlers ^[78–80]^. Second, toddlers’ diffuse and fluctuating attention may have reduced overall stimulus-driven responses ^[81]^. The third reason may have to do with differences in linguistic competence: some of the words and syntactic constructions in *Sesame Street* may be unfamiliar to our toddler participants, which would reduce their ability to build complete linguistic representations of the input. This is the reason that has been put forward to explain the increase in response magnitude for language from early childhood to the late teenage years ^[42]^, as well as in adults learning another language showing stronger responses with increasing proficiency ^[82]^.

### Limitations and future directions

If we want to understand how the human brain comes to comprehend language, we must study toddlers – and yet, this study highlights the obstacles to studying the origins of language in toddlers using fMRI. Two thirds of recruited toddlers who arrived at the fMRI facility did not generate usable data in our paradigm, meaning that our sample – while hard-won – was relatively small and thus not representative of all toddlers in myriad ways. Future work is needed to replicate and extend these results, including using longitudinal study designs to test the stability and changes in language responses across age, conducting more extensive behavioral characterization of language and relating those behavioral measures to brain responses, understanding effects of multilingual exposure, and generalizing to more diverse samples ^[83,84]^. The ability to measure reliable brain responses to language in the earliest stages of linguistic competence may also have clinical applications, by predicting later language challenges.

### Conclusion

In sum, by toddlerhood – when young children are rapidly learning language and beginning to use it communicatively in complex ways – the human brain already shows emergent features of a specialized language system.

## MATERIALS AND METHODS

### Participants

We recruited and attempted to scan 89 toddlers between the ages of 18-36 months (48 female/41 male) over 137 sessions. Most children came in for one visit (n=53), with 26 children coming in for 2 visits, 8 children for 3 visits, and 2 children for 4 visits. The mean age across all visits was 27.15 months (SD= 5.27, range=18-36 months). Average gestational age across all participants was 39.17 weeks (SD=1.54 weeks, range=34-41 weeks, NA=8). Of the 77 participants with reported language exposure, 49 had some regular exposure to language(s) in addition to English, with reported additional languages including: Spanish, Chinese (Mandarin, Cantonese), Korean, Akan, Indonesian, German, Italian, Tamil, Hindi, French, Sinhala/Sinhalese, Vietnamese, Portuguese, Gujarat, Kashmiri, and Arabic. Overall, parental education (averaged across reported caregivers per child) was high (M(SD)=17.95(2.20) years, range=10-23 years, NA=13). See **Table S1** for participant characteristics split by whether participants were included in the final sample. As in the included sample, participants who did not contribute usable fMRI data showed a strong correlation between productive vocabulary and age (*r*=0.77, NA=18; **Fig. S2**). Of the 60 participants not included in analyses, 8 provided some functional data but were excluded for not having at least one usable run, and 52 toddlers did not complete any functional scans (13 of these children began some anatomical scanning).

#### Included participants

29 toddlers (ages 19-36 months, mean(SD)=30.07(4.77) months, 19 female) provided usable functional MRI data and are included in these analyses. 7 of these toddlers include data combined from 2 visits, with the requirement of sessions being within 6 weeks of each other (range: 3-6.57 weeks) for data to be combined. We followed up with parents via email for all participants with usable fMRI data to ask about their child’s handedness (5-37 months after initial participation), with all but two reporting that their child preferred their right hand (1 left, 1 ambidextrous). The average gestational age of included participants was 39.24 weeks (SD=1.46 weeks). This sample included 19 participants identified by caregivers as white, 5 Asian, 1 other, and 4 multiple races selected; 4 identified as Hispanic/Latino. 15 participants had regular exposure to at least one other language in addition to English (sources of additional language varied, such as parents, other family members, and childcare providers). Average parental education was relatively high, ranging from 14 to 22.5 years (mean(SD)=18.46(1.85 years), NA=1).

### Experimental Protocol

The protocol was approved by the MIT Committee on the Use of Humans as Experimental Subjects. Informed consent was provided by parents of participants. Participants were compensated at a rate of $100/session, and $5/survey for parents filling out the MB-CDI ^[52]^.

We employed numerous strategies to acquire fMRI data from toddlers, many inspired by prior research ^[85–88]^. In addition to the stimuli-specific considerations, we also developed various techniques to try with individual children. For instance, we introduced the MRI scanner to children as a “rocket ship,” and framed the visit as a “trip to space.” This framing has been used in a number of pediatric neuroimaging settings, as it provides a fun, plausible explanation for getting into a narrow tube (the “rocket ship”), wearing a head coil (“space helmet”), and hearing background sounds during image acquisition (“noises while blasting through space”). We decorated the scanner with space stickers and used bedsheets with rocket ships.

#### Visit Preparation

Parents were sent detailed instructions before the visit to help them know what to expect. At least one week prior to the visit, parents were asked to start preparing their child by: (1) having their child watch a video we created of another child participating in the study (**Fig. 1A**), (2) reading a book to their child about Elmo going to space in a rocket ship, that outlined the steps they would follow during the visit, (3) playing scanner noises to acclimate their child to the sounds, and (4) practicing listening to favorite songs through in-ear headphones. We told parents that they could bring along (metal-free) favorite stuffed animals and pajamas to help their child feel as comfortable as possible. Parents were also asked to send along favorite videos and songs so that we could play them while getting children set up in the scanner.

#### Visit Protocol

Parents and children met the lead experimenter and the secondary experimenter in the playroom, which contained a mock scanner setup (**Fig. 1A**). While the lead experimenter acquired informed consent from the parent and answered questions, the secondary experimenter played with the child to establish familiarity and rapport. This was important, as the secondary experimenter would be in the scanner room with the child during data acquisition. Next, as much time as needed was taken to get the child comfortable and ready to go into the “real rocket ship.” This could include: reading the “Elmo Goes to Space” book, picking out stuffed animals to come with them into the scanner, practicing putting their stuffed animals in the mock scanner, climbing into the mock scanner themselves, eating a snack, diaper changes, measuring height and weight, taking off shoes, and checking for metal using a handheld metal detector (“magic wand”). Parents changed into metal-free scrubs if they wished to accompany their child in the scanner room, and a second metal detector check for parents and children was completed before entering the scanner control room. To encourage children to enter the scanner room, we let them pick out a space sticker to stick on the “rocket ship.” Then, the goal was to put in the in-ear headphones for hearing protection, secure the headphones in place with an EarBandIt headband, get the child to lay down, place the mirror on top, place the top of the headcoil, then move the child to isocenter in the scanner. Strategies included: playing favorite songs in the headphones (“listen, what song do you hear?”), playing favorite videos on the projected screen at the back of the scanner (“if you lay down and look through this mirror, you can see garbage trucks!”), modeling with stuffed animals (“what does Elmo need next?”), modeling with humans (“it’s mom’s turn!”), and exploring the environment (“let’s press the button to make the bed go up and down”). Oftentimes children would get partway through and fuss out, but we always tried to end on a “success” (e.g., putting in headphones, laying down). In these cases, we always invited parents to come back to try again, and discussed additional strategies to help prepare their child in the interim. If the child did make it into the scanner, as much data were collected as tolerated by the child. The lead experimenter ran the scanner from the control room, using hand signals to communicate with the secondary experimenter, who was in the scanner room monitoring the child and ensuring the hearing protection remained in place.

Parents were either in the scanner room or in the control room. Any time a functional task was not being run (e.g., during transitions and anatomical scans), we played child-friendly videos or videos recommended by the parents. At the end of the scan, children selected a toy to take home and were given a t-shirt.

#### Vocabulary Measure

Parents completed the online version of the English (American) Words & Sentences form of the MB-CDI ^[52]^ within two weeks after the visit. This form is only normed for 16-30 months of age; thus, we only report raw scores. For parents of toddlers with exposure to multiple languages, some if not all reported whether their child knew a given word in any language.

#### fMRI Data Acquisition

fMRI data were acquired from a 3-Tesla Siemens Magnetom Prisma scanner located at the Athinoula A. Martinos Imaging Center at MIT using a 32-channel head coil. For 14 of the included participants, T1-weighted structural images were acquired in 176 interleaved sagittal slices with 1.0mm isotropic voxels (MPRAGE; TA=5:53; TR=2530.0ms; FOV=256mm; GRAPPA parallel imaging, acceleration factor of 2), plus 1 with slightly adjusted parameters (1mm x 1.2mm x 1.2mm due to error in acquisition). Partway through the study, we switched to prioritizing a faster T1 sequence with 1.5mm isotropic voxels for 7 of the included participants (MPRAGE; TA= 2:31; TR=2530.0ms; FOV=192mm; GRAPPA parallel imaging, acceleration factor of 3). Finally, 7 participants did not have a usable T1-weighted structural image, so we used a head scout preliminary image in place of a T1 for alignment (1.6mm isotropic; TA=0:14; TR=3.15ms; FOV=260mm; GRAPPA parallel imaging, acceleration factor of 3; prescan normalized). Functional data were acquired with a gradient-echo EPI sequence sensitive to Blood Oxygenation Level Dependent (BOLD) contrast in 3mm isotropic voxels in 46 interleaved near-axial slices covering the whole brain (EPI factor=70; TR=2s; TE=30.0ms; flip angle=90 degrees; FOV=210mm). fMRI tasks were run from a MacBook Pro laptop and projected onto a 16”x12” screen. Participants viewed the stimuli through a mirror attached to the head coil. Tasks were run through PsychoPy software ^[89]^.

#### fMRI Task

Participants completed up to 4 runs of the fMRI task, which had a 2 × 2 block design with four conditions: forward dialogue, forward monologue, backward dialogue, and backward monologue (**Fig. 1C**; see ^[53]^ for additional details). Stimuli consisted of 20-second edited audiovisual clips from *Sesame Street* during which either two puppets speak to each other (dialogue), or a single puppet addresses the viewer (monologue), with the auditory speech stream played either normally (forward) or reversed so as to be incomprehensible (backward). Dialogue blocks consisted of two characters, both present in the same scene, speaking back-and-forth for a total of 20 seconds, and monologue blocks consisted of two sequential 10-second clips of a character, present alone. In the backward conditions, the audio was reversed for each character rather than across the entire clip, ensuring a continuity of voice-character alignment. For instance, in a backward dialogue block with Elmo and Abby, Elmo’s voice was reversed and played when Elmo was talking, and Abby’s voice was reversed and played while Abby was talking. Each task run contained unique clips, and participants never saw a forward and backward version of the same clip. Each run contained 3 sets of 4 blocks, one of each condition (total of 12 blocks), with 10-second rest periods after each set of 4 blocks and 5-second rest periods after other blocks (5:15 total time per run). Rest periods consisted of clips from the Inscapes video, which was explicitly designed to improve compliance of young children during neuroimaging ^[58]^. These clips involved professionally-produced abstract visuals along with an instrumental soundtrack and were designed as an alternative to resting state scans using a static fixation cross. Forward and backward versions of each clip were counterbalanced between participants (randomly assigned Set A or Set B), but run order was mostly fixed.

### Data Analysis

#### fMRI Preprocessing and Statistical Modeling

FMRI data were first preprocessed using fMRIPrep 24.1.0 ^[90]^, which is based on Nipype 1.8.6 ^[91]^. See **Supplementary Text** for full preprocessing pipeline details. We manually reviewed preprocessing outputs from each participant before proceeding with data analysis. We used a lab-specific script that uses Nipype to combine tools from several different software packages for first-level modeling. Each event regressor was defined as a boxcar convolved with a standard double-gamma HRF, and a high-pass filter (1/210 Hz) was applied to both the data and the model. Artifact detection was performed using Nipype’s RapidART toolbox (an implementation of SPM’s ART toolbox).

Individual TRs were marked as outliers if (1) there was more than 1 unit of frame displacement (see **Supplementary Text** and **Figs. S6-S7** for more lenient and stricter outlier thresholds), or (2) the average signal intensity of that volume was more than 3 standard deviations away from the mean average signal intensity. We included one regressor per outlier volume. In addition, we included a summary movement regressor (framewise displacement) and 6 anatomical CompCor regressors to control for the average signal in white matter and CSF. We applied a 6mm smoothing kernel to preprocessed BOLD images. The first-level model was run using FSL’s GLM in MNI space. Subject level modeling was performed with in-lab scripts using Nipype. Specifically, FSL’s fixed effects flow was used to combine runs at the level of individual participants. A subject level model was created for each set of usable runs per contrast for the task (up to 4 runs). Runs with more than 35% of timepoints marked as outliers and runs with less than 3:20 of data collected (fewer than 2 blocks per condition) were excluded from analysis. Output average magnitudes in each voxel in the second level model were then passed to the group level model.

#### Whole Brain Analysis

Group modeling used in-lab scripts that implemented FSL’s RANDOMISE to perform a nonparametric one-sample t-test of the contrast values against 0 (5000 permutations, MNI space). We used Threshold-Free Cluster Enhancement (TFCE; ^[92]^) for multiple comparisons correction with variance smoothing (6mm kernel). Results were thresholded at corrected p < 0.05. In toddlers, we show the uncorrected group random effects analyses at a lenient threshold (T>1.70, for a one-sample T-test with 28 degrees of freedom at p<0.05), since there was no significant activation at the corrected threshold. For language (Language>Control), we used the [forward dialogue + forward monologue] > [backward dialogue + backward monologue] contrast. For social processing (Dialogue>Monologue), we used the [forward dialogue + backward dialogue] > [forward monologue + backward monologue] contrast.

#### Functional Region of Interest (fROI) Analysis

17 participants had more than one usable run of the functional task and were included in fROI analyses. We iteratively defined subject-specific functional regions of interest (fROIs) for language as the top 100 voxels activated in an individual, within each of five predefined language search spaces, for the Forward>Backward contrast in n-1 runs, where n is the total usable runs ^[54]^. The five language search spaces in the left hemisphere included: Left IFGorb, Left IFG, Left MFG, Left AntTemp, and Left PostTemp (similar to Fedorenko et al, 2010; in this case, 5 parcels in left hemisphere were created based on a group-level probabilistic activation overlap map for a sentences>nonwords contrast in 220 adult participants; parcels downloaded from https://evlab.mit.edu/funcloc/). We also looked within the mirror of these search spaces in the right hemisphere (i.e., right hemisphere language homotopes). We used an in-lab script that iteratively used the z-stat image of each n-1/n combined runs (i.e., each ‘fold’) to determine the top 100 voxels for a given subject and ROI. We then used the cope image from the left-out run of a given iteration to extract the betas from these selected top voxels, then averaged these betas together per participant. In an exploratory analysis, we also used the top 10% voxels per parcel to define fROIs, which did not affect the results (**Table S9**).

#### Lateralization

Another common measure of language in the brain is the laterality index; that is, how many more voxels are activated for language on the left compared to the right. For all participants, we calculated the laterality index using the formula, LI=(Vleft-Vright)/(Vleft+Vright), where V refers to the number of suprathreshold voxels for Language>Control ^[37,93]^. For exploratory analyses, we calculated LI using a threshold of z>1.64 and a cluster threshold of k>=10 voxels, constrained to the left language search spaces and their mirrored right hemisphere homotope search spaces. This is a more lenient threshold than we preregistered, given the small number of suprathreshold voxels at the preregistered level of p<0.01. To ensure that the threshold selected did not impact results, we also conducted LI analyses with thresholds of z>0.84, z>1.28, z>1.96, z>2.33, and z>2.58 (**Table S8**). In an exploratory analysis, we also calculated LI for the Dialogue>Monologue contrast, using the same approach (threshold z>1.64, cluster threshold k>=10 voxels).

#### Univariate Analyses

Statistical analyses were conducted in R (Version 2022.07.1; R Core Team, 2022), using the average activation per condition within fROIs. Conditions were compared using linear mixed effects models; t-tests used Satterthwaite’s method. To test for network-level fixed effects, with region of interest (ROI) and participants modeled as random effects, we used: lmer(mean_topvoxels_extracted ∼f_or_b * d_or_m * t_or_f + age + (1|ROI) + (1|participantID), REML = FALSE), where f_or_b is forward or backward (coded 1, −1, respectively), d_or_m is dialogue or monologue (coded 1, −1), t_or_f is temporal or frontal (1, - 1, respectively), age is in months, and ROI is region of interest within the network. Significance was determined at a level of p < 0.05. To test for interactions within individual regions, we used: lmer(mean_topvoxels_extracted ∼f_or_b * d_or_m + age + (1|participantID), REML = FALSE). Significance was determined at a level of p < 0.05 Bonferroni corrected for the number of ROIs (five for canonical language regions and five for right-hemisphere language regions; p < 0.01).

#### Adult Comparison Group

20 adults (ages 18-30 years, mean(SD)= 23.85(3.70)) previously participated in a study using a similar fMRI task ^[53]^. Adults completed 4 runs of the functional task, which differed from the toddler version in the following ways: (1) a black screen with a fixation cross was used for all baseline periods, rather than *Inscapes* clips ^[58]^, (2) adults responded to an attention check with an in-scanner button box following each block (i.e., press a button when a still image of Elmo appeared on the screen), and (3) the longer fixation blocks were 22 seconds (we reduced this to 10 seconds for the toddler version).

## Acknowledgments

We are grateful to the exceptional team of undergraduate students that assisted with various aspects of this project: Somaia Saba, Hana Ro, Rebecca O’Connor, Bianca Santi, Sofia Riskin, Matthew Soza, Maria Garcia-Garcia, Linette Kunin, Michelle Hung, Maria Gabas, and Laila Salgado. We also thank Kirsten Lydic for technical support, Kris Brewer from the Center for Brains, Minds, and Machines for filming and editing the toddler preparation video, and Steve Shannon and Atsushi Takahashi from the Athinoula A. Martinos Imaging Center at MIT for supporting toddler scanning. Finally, thank you to the participating families for making this research possible.

