## Supplement for "The emergence of the language system in the toddler brain"

### SUPPLEMENTARY MATERIALS

##### Behavioral Task Validation in Toddlers

To verify that toddlers would attend to all task conditions, we conducted an online behavioral pilot study prior to fMRI data collection. 36 participants (ages 14-34 months, mean(SD)=25.06(4.52) months) watched the same 20-second video clips of *Sesame Street* that ended up being used for the fMRI task (forward monologue, backward monologue, forward dialogue, and backward dialogue). Forward and backward versions of each clip were counterbalanced between participants, and trials were presented in a balanced, pseudorandom order with the order of trial conditions counterbalanced between participants, such that each version began with a different condition. There were a total of 12 trials per condition (48 total trials). Children continued the experiment until they watched all 48 trials or decided to stop (approximately 17 minutes total). Data collection was conducted over Zoom with a live experimenter. The session was recorded and stored securely for offline coding. The caregiver was instructed not to direct the child's attention back to the screen during the videos, and the caregiver and child were told to let the experimenter know if the child was "all done." The experimenter also sometimes ended the experiment if the child seemed upset or was no longer paying any attention to the experiment. The child was instructed to look at the screen at the beginning of the experiment (e.g., "Look, CHILD! Here we go!"). An attention-getter appeared on the screen in the four corners to help experimenters determine where the edges of the child's screen were for coding of looking behavior. An attention getter appeared after every trial in the center of the screen. Looking time was coded offline using DataVyu [1]. A member of the research team first identified the timestamp for the start of each trial based on the end of the attention getter. Looks were coded as “on” or “off” depending on whether the child was looking at the screen. For each child, we calculated the average looking time for all the trials they watched. Participants watched 3-48 valid trials (mean(SD)=29.58(13.24) trials) out of a possible 48 trials. A one-way repeated measures ANOVA was conducted to examine the effect of condition on average looking time for all valid trials per condition per participant; looking time did not differ between conditions (**Fig. S1**; $F(3, 102)=2.109$, $p=.10$). Furthermore, for the trials children watched, average looking time per condition per child was close to the 20s trial duration (range=7.4-20.3s, mean(SD)=16.8(3.1) s).

#### fMRI Task Validation in Adults

To validate the experimental task, we conducted an fMRI study with 20 adult participants (see [2] for full study details). To determine whether our *Sesame Street* task recruited canonical language regions for the Forward>Backward contrast, we compared responses to the Intact>Degraded contrast from an auditory language localizer [3]. First, across all subjects, we quantified the overlap at the whole-brain level using the group random effects analysis results. Specifically, we calculated Dice coefficients of similarity to capture the extent of overlap in thresholded activation maps, using the formula: Dice coefficient = $2*V_{overlap}/(V_1+V_2)$, where $V_{overlap}$ refers to the number of supra-threshold voxels identified in both tasks, $V_1$ and $V_2$ refer to the number of supra-threshold voxels for each of the two tasks, respectively [4]. We used a threshold of $p<0.001$ (TFCE corrected) and cluster threshold of $k>=10$ voxels. Dice coefficients can be described as: low (0-0.19), low-moderate (0.2-0.39), moderate (0.4-0.59), high-moderate (0.6-0.79), and high (0.8-1) [4]. We also calculated the overlap at the whole brain level for each individual subject using a threshold of $p<0.001$. Just as we expected overlap across the whole brain, we also expected a high degree of overlap between the Forward>Backward and Intact>Degraded contrasts within each language region, per subject. We calculated and report Dice coefficients to capture the extent of overlap within each language search space, using a threshold of $p<0.001$ and cluster threshold of $k>= 10$ voxels to identify suprathreshold voxels. Finally, we calculated the overlap in the top 100 language-selective voxels (e.g., our definition of an fROI) in each subject using the LIT task and using the auditory language localizer task, as a measure of overlap in how these tasks would define language fROIs. The contrast of Forward versus Backward speech in the *Sesame Street* clips robustly activated the same cortical regions as the contrast of Intact versus Degraded speech using the classic auditory language localizer, both at the group level (Dice coefficient=0.71; high-moderate overlap) and at the level of individual subjects, across the whole brain and within language parcels. The mean overlap for all participants was classified as high in left AntTemp and PostTemp regions, and high-moderate in left IFGorb, IFG, and MFG.

#### Preregistration

The key hypotheses and analysis plan for this study were preregistered on OSF, including our sample size of 16 toddlers with at least two usable runs of the fMRI task. We deviated from the preregistration in a few ways. First, we used an updated version of fMRIprep for final analyses (version 24.1.0). Second, for run exclusion, our preregistered criteria (i.e., runs that did “not include usable data from all 4 conditions”) was underspecified, so we updated our inclusion criteria to specify that a run must be at least 3:20 to be included (i.e., 2 blocks per condition presented). Third, we decided to included runs with less than 35% outliers (rather than the preregistered threshold of 33% outliers). Finally, we used less stringent thresholds for group random effects analyses (uncorrected $T>1.70$) and individual lateralization analyses ($Z>1.64$) because toddler brain responses did not reach the stringent preregistered thresholds.

#### Preprocessing Details

The below boilerplate text was automatically generated by fMRIPrep with the express intention that users should copy and paste this text into their manuscripts unchanged. It is released under the CC0 license.

Results included in this manuscript come from preprocessing performed using fMRIPrep 24.1.0 ([5,6]; RRID:SCR_016216), which is based on Nipype 1.8.6 ([7,8]; RRID:SCR_002502).

**Anatomical data preprocessing.** A total of 1 T1-weighted (T1w) images were found within the input BIDS dataset. The T1w image was corrected for intensity non-uniformity (INU) with N4BiasFieldCorrection [9], distributed with ANTs 2.5.3 ([10]; RRID:SCR_004757), and used as T1w-reference throughout the workflow. The T1w-reference was then skull-stripped with a Nipype implementation of the antsBrainExtraction.sh workflow (from ANTs), using OASIS30ANTs as target template. Brain tissue segmentation of cerebrospinal fluid (CSF), white-matter (WM) and gray-matter (GM) was performed on the brain-extracted T1w using fast (FSL (version unknown), RRID:SCR_002823, [11]). Brain surfaces were reconstructed using recon-all (FreeSurfer 7.3.2, RRID:SCR_001847, [12]), and the brain mask estimated previously was refined with a custom variation of the method to reconcile ANTs-derived and FreeSurfer-derived segmentations of the cortical gray-matter of Mindboggle (RRID:SCR_002438, [13]). Volume-based spatial normalization to two standard spaces (MNI152NLin6Asym, MNI152NLin2009cAsym) was performed through nonlinear registration with antsRegistration (ANTs 2.5.3), using brain-extracted versions of both T1w reference and the T1w template. The following templates were were selected for spatial normalization and accessed with TemplateFlow (24.2.0, [14]): FSL's MNI ICBM 152 non-linear 6th Generation Asymmetric Average Brain Stereotaxic Registration Model [[15], RRID:SCR_002823; TemplateFlow ID: MNI152NLin6Asym], ICBM 152 Nonlinear Asymmetrical template version 2009c [[16], RRID:SCR_008796; TemplateFlow ID: MNI152NLin2009cAsym]. Grayordinate "dscalar" files containing 91k samples were resampled onto fsLR using the Connectome Workbench [17].

**Functional data preprocessing.** For each of the BOLD runs found per subject (across all tasks and sessions), the following preprocessing was performed. First, a reference volume was generated, using a custom methodology of fMRIPrep, for use in head motion correction. Head-motion parameters with respect to the BOLD reference (transformation matrices, and six corresponding rotation and translation parameters) are estimated before any spatiotemporal filtering using mcflirt (FSL , [18]). The BOLD reference was then co-registered to the T1w reference using bbregister (FreeSurfer) which implements boundary-based registration [19]. Co-registration was configured with six degrees of freedom. Several confounding time-series were calculated based on the preprocessed BOLD: framewise displacement (FD), DVARS and three region-wise global signals. FD was computed using two formulations following Power (absolute sum of relative motions, [20]) and Jenkinson (relative root mean square displacement between affines, [18]). FD and DVARS are calculated for each functional run, both using their implementations in Nipype (following the definitions by [20]). The three global signals are extracted within the CSF, the WM, and the whole-brain masks. Additionally, a set of physiological regressors were extracted to allow for component-based noise correction (CompCor, [21]). Principal components are estimated after high-pass filtering the preprocessed BOLD time-series (using a discrete cosine filter with 128s cut-off) for the two CompCor variants: temporal (tCompCor) and anatomical (aCompCor). tCompCor components are then calculated from the top 2% variable voxels within the brain mask. For aCompCor, three probabilistic masks (CSF, WM and combined CSF+WM) are generated in anatomical space. The implementation differs from that of Behzadi et al. in that instead of eroding the masks by 2 pixels on BOLD space, a mask of pixels that likely contain a volume fraction of GM is subtracted from the aCompCor masks. This mask is obtained by dilating a GM mask extracted from the FreeSurfer's aseg segmentation, and it ensures components are not extracted from voxels containing a minimal fraction of GM. Finally, these masks are resampled into BOLD space and binarized by thresholding at 0.99 (as in the original implementation). Components are also calculated separately within the WM and CSF masks. For each CompCor decomposition, the k components with the largest singular values are retained, such that the retained components' time series are sufficient to explain 50 percent of variance across the nuisance mask (CSF, WM, combined, or temporal). The remaining components are dropped from consideration. The head-motion estimates calculated in the correction step were also placed within the corresponding confounds file. The confound time series derived from head motion estimates and global signals were expanded with the inclusion of temporal derivatives and quadratic terms for each [22]. Frames that exceeded a threshold of 0.5 mm FD or 1.5 standardized DVARS were annotated as motion outliers. Additional nuisance timeseries are calculated by means of principal components analysis of the signal found within a thin band (crown) of voxels around the edge of the brain, as proposed by [23]. The BOLD time-series were resampled onto the left/right-symmetric template "fsLR" using the Connectome Workbench [17]. Grayordinates files [17] containing 91k samples were also generated with surface data transformed directly to fsLR space and subcortical data transformed to 2 mm resolution MNI152NLin6Asym space. All resamplings can be performed with a single interpolation step by composing all the pertinent transformations (i.e. head-motion transform matrices, susceptibility distortion correction when available, and co-registrations to anatomical and output spaces). Gridded (volumetric) resamplings were performed using nitransforms, configured with cubic B-spline interpolation.

Many internal operations of fMRIPrep use Nilearn 0.10.4 ([24], RRID:SCR_001362), mostly within the functional processing workflow. For more details of the pipeline, see the section corresponding to workflows in fMRIPrep's documentation.

##### Robustness check: Toddler template

In addition to using a standard adult MNI template brain, we also registered the data to a toddler template during preprocessing ([16]; downloaded from: github.com/templateflow/tpl-MNIInfant; MNIInfant: cohort 9, 27-33 months). The registration failed for one toddler participant, so we ran analyses for N=28 toddlers with usable data (N=16 with multiple usable fMRI runs) using the same motion cutoffs as the main analyses (>1mm FD as a motion outlier and >35% outliers to exclude a run). For fROI analyses, language parcels were transformed into the toddler template space using FSL's antsApplyTransforms [25]. Results were similar to the analyses with the standard adult MNI template (see **Fig. S5**).

Using the same linear mixed effects model to examine the effects of language comprehensibility (forward vs. backward speech), social context (dialogue vs. monologue), and cortical lobe (temporal lobe vs. frontal lobe) on univariate responses in individually-defined language regions in toddlers with at least two usable runs (N=16), we found that the canonical left-hemisphere language regions showed higher responses to language than the control conditions in toddlers (Forward>Backward: Est= 0.213, S.E.= 0.045, t-value= 4.702, p-value<0.001). These regions did not show sensitivity to social context (Dialogue>Monologue: Est= 0.017, S.E.= 0.045, t-value= 0.371, p-value= 0.711). Temporal regions showed larger responses to all stimuli than frontal regions (Temporal>Frontal: Est= 0.144, S.E.= 0.045, t-value=3.186, p-value=0.002), but there was no interaction between cortical lobe and either functional contrast. Age was also not significant in the model (Est.= 0.010, S.E.= 0.025, t-value=0.426; p-value=0.676). In individual language regions of interest, a significantly stronger response to language than the control conditions was present in left IFGorb (Forward>Backward: Est=0.321, S.E.= 0.090, t-value=3.582, p=0.0008, uncorrected), left IFG (Forward>Backward: Est=0.278, S.E.= 0.099, t-value=2.812, p=0.007, uncorrected), and left AntTemp (Forward>Backward: Est=0.226, S.E.= 0.055, t-value=4.087, p=0.0002, uncorrected), and a marginally stronger response to language was present in left PostTemp (Forward>Backward: Est=0.160, S.E.= 0.088, t-value=1.806, p=0.077, uncorrected). The language contrast was not significant in left MFG. Just as in the model across the left hemisphere language network, Dialogue>Monologue, interactions, and age were not significant in individual regions' models.

We again calculated a laterality index (LI) for language by comparing the proportion of voxels significantly activated by language (Language>Control contrast) in the left hemisphere canonical language parcels and the mirrored right hemisphere homotope parcels in each participant (threshold: Z>1.64, cluster threshold k>=10). Overall, LI was significantly greater than 0, confirming that language responses were left-lateralized in toddlers (mean(SD)= 0.255(0.69); one-sample t-test for LI>0: t= 1.95, p-value= 0.031). LI was not correlated with age in toddlers (*r*= 0.193; p= 0.162). We also calculated LI for the social processing Dialogue>Monologue contrast within the same parcels. The Dialogue>Monologue response was right-lateralized overall (mean(SD)= -2.119(0.64); one-sample t-test for LI<0: t= -2.119, p-value= 0.022), and this response became more right-lateralized with age (*r*= -0.566; p= 0.001).

#### Robustness check: Varying motion tolerance

To understand the effects of different approaches to handling motion, we applied more lenient (infant-based) and stricter (adult-based) criteria for motion artifact identification and data inclusion.

**Infant-based thresholds**. First, we applied criteria that are similar to some previous fMRI studies in awake infants [26]. We defined motion outliers as >3mm FD and excluded runs with >50% outliers. Applying these standards to our dataset, we had 21 toddlers with 2 or more usable runs, and 10 toddlers with 1 usable run of fMRI data. Results were qualitatively similar to the main analyses, replicating the overall findings (see **Fig. S6**).

Using the same linear mixed effects model to examine the effects of language comprehensibility (forward vs. backward speech), social context (dialogue vs. monologue), and cortical lobe (temporal lobe vs. frontal lobe) on univariate responses in individually-defined language regions in toddlers with at least two usable runs (N=21), we found that the canonical left-hemisphere language regions showed higher responses to language than the control conditions in toddlers (Forward>Backward: Est= 0.269, S.E.= 0.054, t-value= 4.954, p-value<0.001). These regions did not show sensitivity to social context (Dialogue>Monologue: Est= -0.003, S.E.= 0.054, t-value= -0.058, p-value= 0.953). Temporal regions showed larger responses to all stimuli than frontal regions (Temporal>Frontal: Est= 0.120, S.E.= 0.054, t-value= 2.210, p-value= 0.028), but there was no interaction between cortical lobe and either functional contrast. Age was also not significant in the model (Est.= 0.012, S.E.= 0.021, t-value= 0.587; p-value= 0.564). In individual language regions of interest, a significantly stronger response to language than the control conditions was present in left IFGorb (Forward>Backward: Est=0.329, S.E.= 0.107, t-value=3.065, p=0.003, uncorrected), left IFG (Forward>Backward: Est=0.352, S.E.= 0.124, t-value=2.841, p=0.006, uncorrected), and left AntTemp (Forward>Backward: Est=0.307, S.E.= 0.103, t-value=2.977, p=0.004, uncorrected), and a marginally stronger response to language was present in left PostTemp (Forward>Backward: Est=0.217, S.E.= 0.117, t-value=1.852, p=0.0687, uncorrected). The language contrast was not significant in left MFG. Just as in the model across the left hemisphere language network, Dialogue>Monologue, interactions, and age were not significant in individual regions' models.

We again calculated a laterality index (LI) for language by comparing the proportion of voxels significantly activated by language (Language>Control contrast) in the left hemisphere canonical language parcels and the mirrored right hemisphere homotope parcels in each participant (threshold: Z>1.64, cluster threshold k>=10). Overall, LI was significantly greater than 0, confirming that language responses were left-lateralized in toddlers (mean(SD)= 0.247(0.628); one-sample t-test for LI>0: t=2.190, p-value=0.018). LI was not correlated with age in toddlers (*r*=0.179; p=0.168). We also calculated LI for the social processing Dialogue>Monologue contrast within the same parcels. The Dialogue>Monologue response was not right-lateralized overall (mean(SD)= -0.136(0.573); one-sample t-test for LI<0: t= -1.304, p-value= 0.101), and there was a marginal effect of becoming more right-lateralized with age (*r*= -0.290; p= 0.060).

**Adult-based thresholds**: Next, we applied the same criteria as in our prior study in adults [2]. We defined motion outliers as >0.4mm FD and excluded runs with >25% outliers. Applying these standards to our dataset, we were left with 7 toddlers with 2 or more usable runs, and 9 toddlers with 1 usable run of fMRI data. Results differed qualitatively from the main analyses and the exploratory analyses at the more lenient infant-based thresholds (**Fig. S7)**. With only 7 toddlers that had 2 or more usable runs, we did not run the linear mixed effect models, but qualitatively, we did not observe a language-specific response in functionally-defined regions of interest. The whole brain random effects analysis likewise showed little difference between the language and control conditions. Laterality index (LI) for language (Language>Control; threshold: Z>1.64, cluster threshold k>=10) was not significantly greater than 0 (mean(SD)= 0.158(0.714); one-sample t-test for LI>0: t= 0.886, p-value= 0.195) and was not correlated with age (*r*= 0.226; p= 0.201). Laterality index (LI) for the social processing contrast (Dialogue>Monologue; threshold: Z>1.64, cluster threshold k>=10) was not significantly less than 0 (mean(SD)= -0.086(0.552); one-sample t-test for LI<0: t= -0.623, p-value= 0.271), but responses did become more right lateralized with age (*r*= -0.497; p= 0.025). Overall, these results support the general pattern we see across all analyses: individual toddlers are noisy, and this noise is not just due to using more lenient motion thresholds.

##### Robustness check: Varying fROI definition (top 10%)

In addition to our pre-registered approach of selecting the top 100 voxels in each language parcel, we performed an exploratory analysis selecting the top 10% of voxels responding to the Language>Control contrast in each parcel. We again used a linear mixed effects model to examine the effects of comprehensibility (forward vs. backward speech), social context (dialogue vs. monologue), and cortical area (temporal vs. frontal) on univariate responses in individually-defined language regions (this time, using the top 10% of voxels) while controlling for age. The set of canonical left-hemisphere language regions showed higher responses to language than the control conditions in toddlers (Forward>Backward: Est=0.174, S.E.=0.050, t-value=3.466, p-value=0.0006). There was again no effect of social context (Dialogue>Monologue: Est=0.089, S.E.=0.050, t-value=1.773, p-value=0.077), and there was a main effect of cortical area (Temporal>Frontal: Est=0.139, S.E.=0.052, t-value=2.692, p-value=0.048). Age was not significant in the model (Est=0.012, S.E.= 0.022, t-value=0.519, p-value=0.610). Responses in individual language parcels are shown in **Table S9**.

##### Robustness check: Reliability across runs for lateralization analyses

While toddlers, like adults, showed overall left-lateralization for language across the group, their responses were variable across individuals (see **Fig. 4**). To understand whether this variability reflected true individual differences or noisy data, we examined LI for individual runs in toddlers with multiple usable runs (N=17). LI was calculated for each run (rather than each participant) as in the primary analyses (Language>Control; threshold: Z>1.64, cluster threshold k>=10). We then used an intra-class correlation analysis to estimate how much of the total variance in LI is due to differences between participants vs. total variance, using the psych::ICC package in R. ICC values were 0 and not statistically significant (p=0.47), meaning that there is no evidence of consistent LI scores across runs.

##### Right hemisphere homotopes of language regions

We tested whether right hemisphere homotopes of the language network are language-selective in toddlers. The right hemisphere homotope language regions did show a main effect of cortical lobe (Temporal>Frontal: Est = 0.160, S.E. = 0.050, t = 3.195, p = 0.0015). However, when we conducted an exploratory linear mixed effects model with both left and right regions included (model: AverageBeta~ForwardvsBackward * LeftvsRight + Age + (1|participant) + (1|ROI)), we did not find a significant interaction between language comprehension and hemisphere (Forward>Backward * Left>Right: Est = 0.063, S.E. = 0.037, t = 1.713, p = 0.087), though there was a main effect of language comprehension (Forward>Backward: Est = 0.115, S.E. = 0.037, t = 3.133, p = 0.002). When examining individual regions, only right AntTemp showed a marginally stronger response to language than the control conditions (Forward>Backward: Est = 0.193, S.E. = 0.090, t = 2.140, p = 0.037, uncorrected; see Table S5 for full model results for individual regions).

##### Exploratory: Impacts of language experience

To check whether productive vocabulary or exposure to languages other than English impacted brain responses, we performed exploratory analyses controlling for multi-language exposure and CDI scores in the linear mixed effects model that examined the effects of language comprehensibility (forward vs. backward speech), social context (dialogue vs. monologue), and cortical lobe (temporal lobe vs. frontal lobe) on univariate responses in individually-defined language regions in toddlers with at least two usable runs (N=17).

AverageBeta~ForwardvsBackward *DialoguevsMonologue * TemporalvsFrontal + Age + **MultLang** + **CDI** + (1|participant) + (1|ROI)

When only multi-language exposure was included in the model (not CDI scores), the results were similar to the main analyses. There was still a main effect of comprehensibility in left-hemisphere language regions (Forward>Backward: Est.= 0.175, S.E.= 0.050, t-value= 3.492, p-value= 0.0005) and a main effect of cortical lobe (Temporal>Frontal: Est.= 0.162, S.E.= 0.060, t-value= 2.703, p-value= 0.047), but no significant effect of social context (Dialogue>Monologue: Est.= 0.093, S.E.= 0.050, t-value= 1.869, p-value= 0.062). Multi-language exposure, age, and interactions were nonsignficant (all ps> 0.37).

When only CDI scores were included in the model (not multi-language exposure), the results slightly differed from the main analyses. Note that, because CDI scores were missing from 3 participants, this analysis only included 14 toddlers rather than 17. There was still a main effect of comprehensibility in left-hemisphere language regions (Forward>Backward: Est.= 0.129, S.E.=0.050, t-value= 2.576, p-value= 0.011), and in this case there was an effect of social context (Dialogue>Monologue: Est.=0.105, S.E.=0.050, t-value= 2.099, p-value= 0.037) but no effect of cortical lobe (Temporal>Frontal: Est.=0.138, S.E.=0.058, t-value= 2.391, p-value= 0.068). Multi-language exposure, age, and interactions were nonsignficant (all ps> 0.34).

With both multi-language exposure and CDI scores in the model (as above, limiting to N=14 due to missing CDI scores from 3 participants), there was still an effect of comprehensibility in left-hemisphere language regions (Forward>Backward: Est.=0.129, S.E.=0.050, t-value=2.576, p-value=0.0106). In this case, there was an effect of social context (Dialogue>Monologue: Est.=0.105, S.E.= 0.050, t-value=2.099, p-value=0.037) and no effect of cortical lobe (Temporal>Frontal: Est.=0.138, S.E.=0.058, t-value=2.393, p-value=0.067). Age, multi-language exposure, CDI scores, and interactions were nonsignficant (all ps>0.29).

We also conducted a Welch two-sample t-test to ensure that lateralization for language did not differ depending on multi-language exposure, and the groups did not differ (N=15 with multi-language exposure, mean(SD)= 0.315(0.618); N=14 without multi-language exposure, mean(SD)= 0.198(0.619); two-sample t-test: t-value= 0.509, p-value= 0.615).

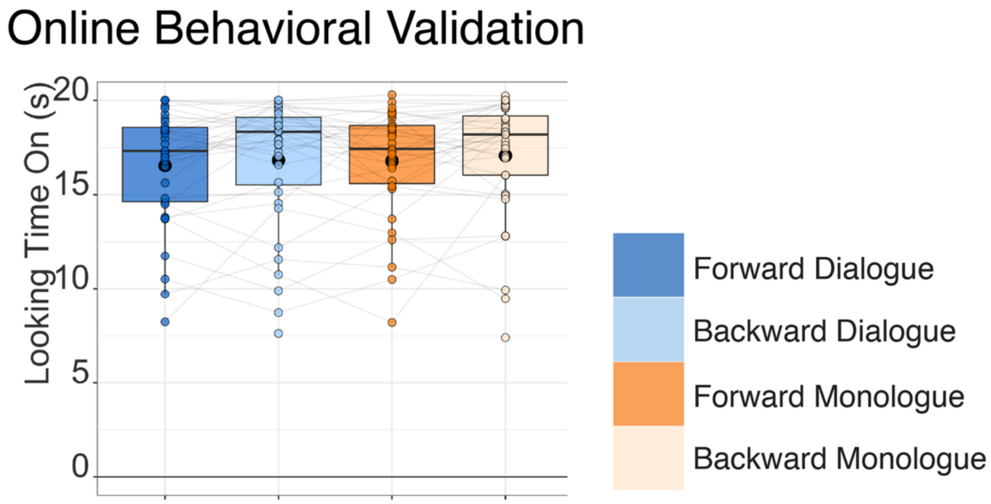

**Fig. S1.**
Toddlers (N=36, aged 14-34 months; not included in the fMRI study) viewed the *Sesame Street* stimuli in a passive-viewing online pilot study. Videos were coded based on whether the child was looking at the screen during video presentation, for each 20-second trial. Individual dots show each participant's looking time (in seconds) averaged across all valid trials per condition; light gray lines connect the same participant; large black dot shows mean. There were no significant differences in looking time between conditions.

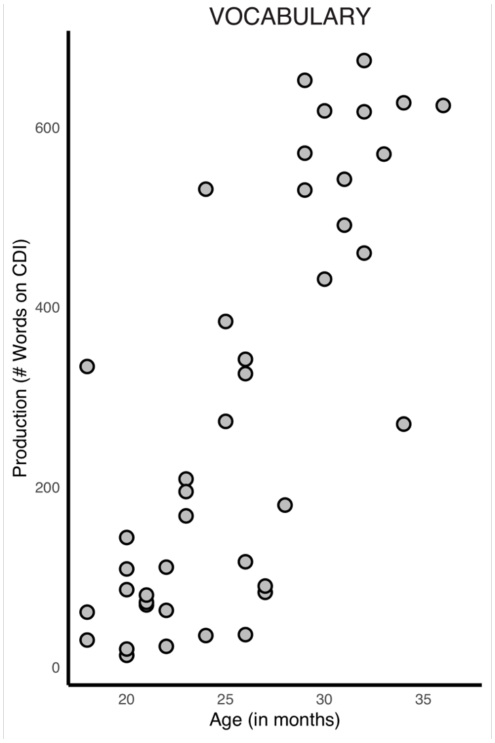

**Fig. S2.**
Age in months and raw score for the CDI from the first time the caregiver filled out the CDI, for children without usable fMRI data (plot includes N=42 with CDI data).

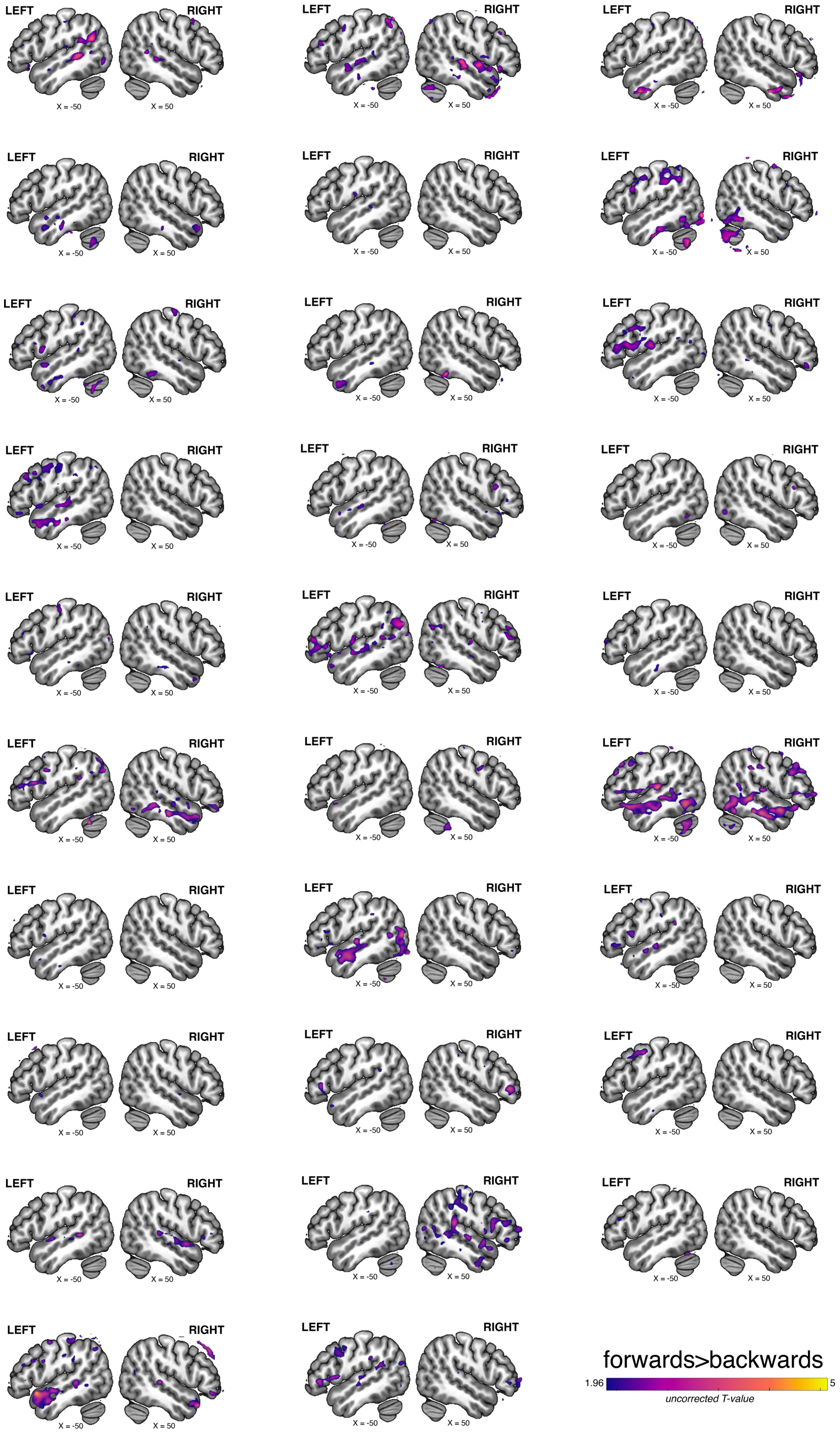

**Fig. S3.**

Responses for the Language>Control contrast are visualized for individual toddler participants at an uncorrected threshold T>1.96.

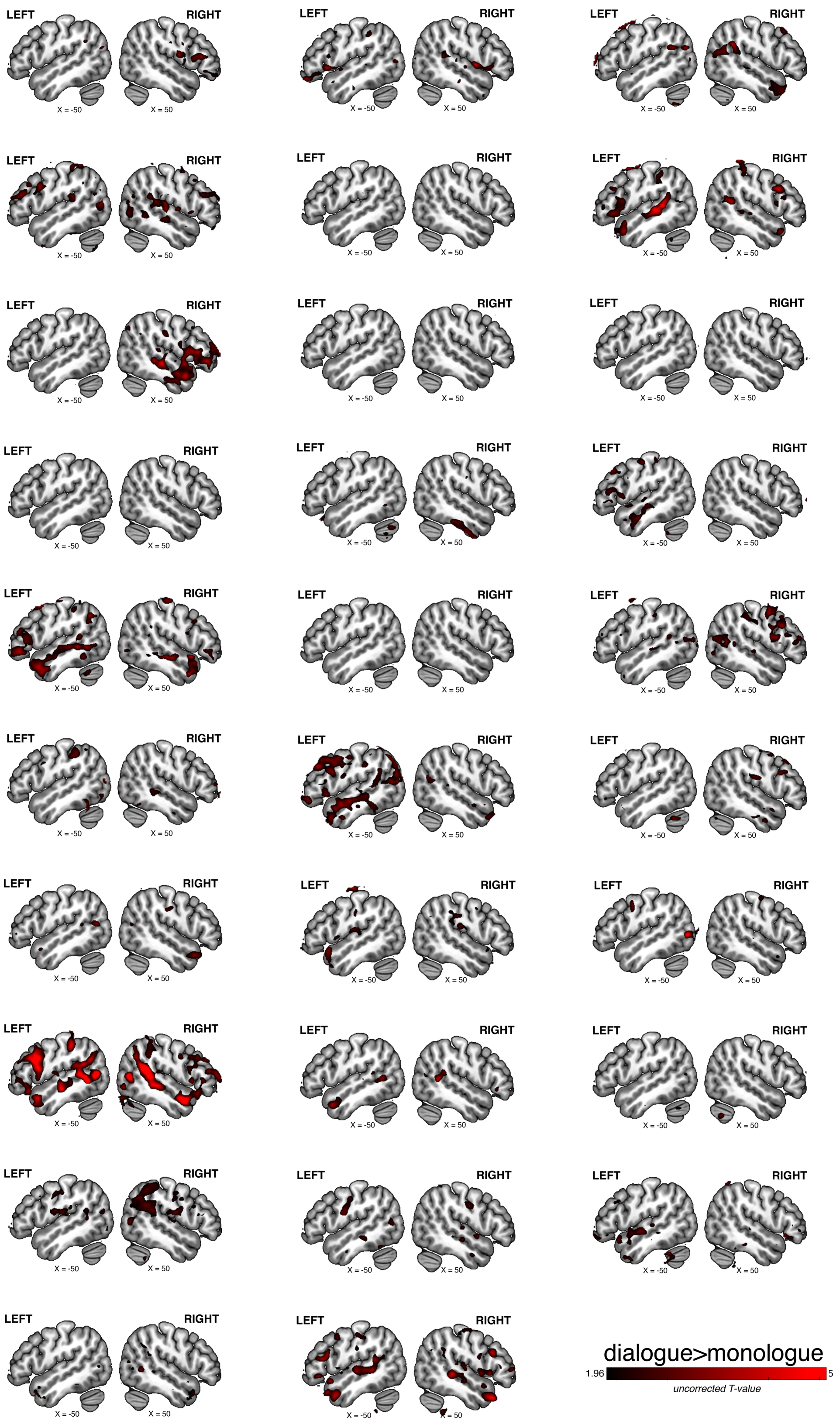

**Fig. S4.**

Responses for the Dialogue>Monologue contrast are visualized for individual toddler participants at an uncorrected threshold T>1.96.

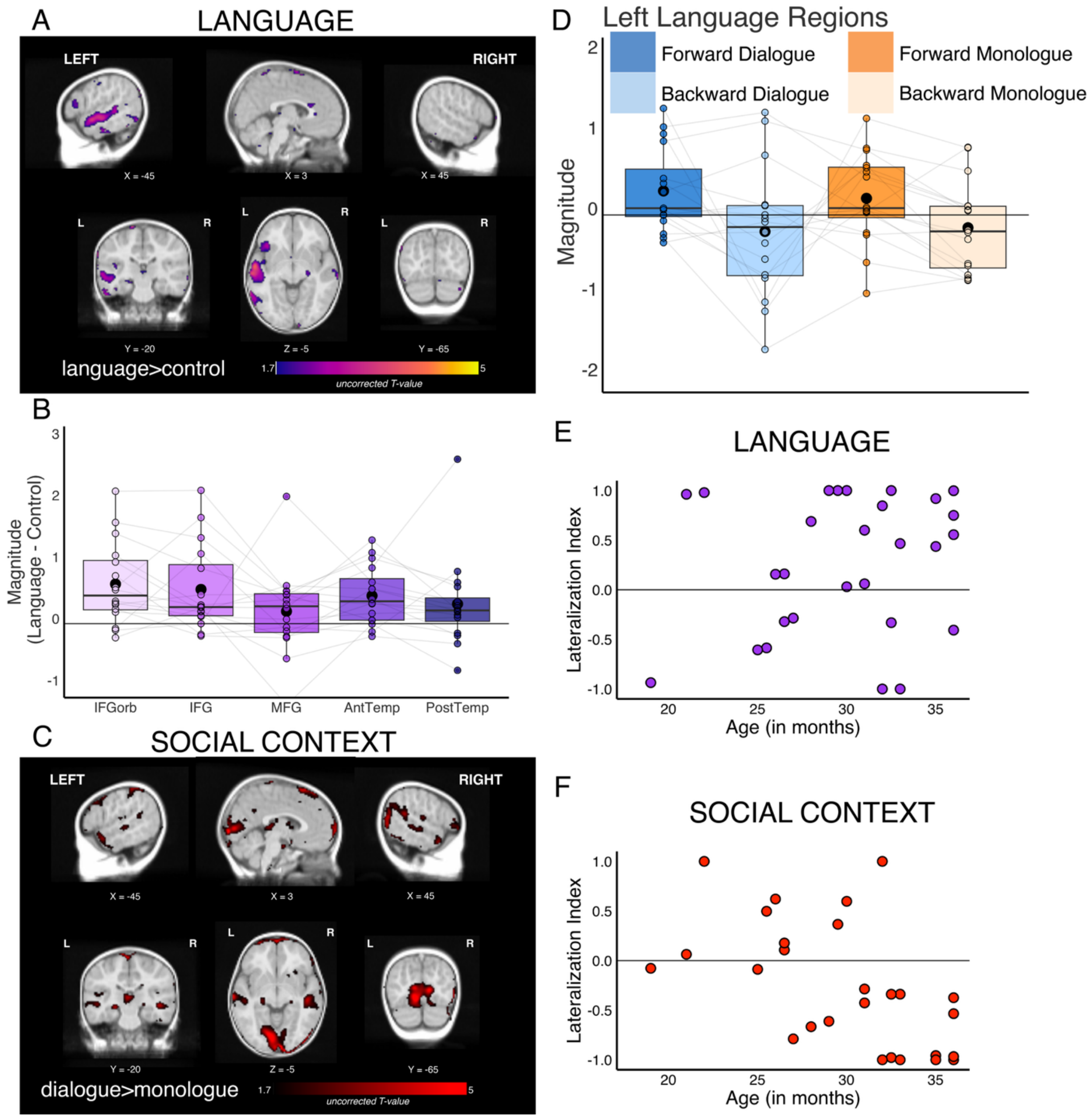

**Fig. S5.**

Results using a toddler template (MNIInfant: cohort 9, 27-33 months, [16]): we defined motion outliers as >1mm FD and excluded runs with >35% outliers (consistent with main analyses). A group random effects analyses was conducted in the full sample of toddlers (N=28; note that one toddler failed preprocessing quality control) with usable data at this threshold for **(A)** the language contrast and **(C)** the social context contrast. Toddler group responses are visualized at an uncorrected threshold T>1.70. **(D)** For participants with multiple usable functional runs (N=16), fROIs for the top 100 voxels were iteratively identified using the Language>Control contrast in each language parcel using a leave-one-run-out approach. Boxplots show response magnitude for held-out runs (independent data) for each condition averaged across all regions. **(B)** For each language region, we plot the difference in magnitude between language and control conditions in each individually-defined region (purple). Individual dots show each participant's response; light gray lines connect the same participant; large black dot shows mean. **(E)** For all 28 toddlers with usable data, lateralization index was calculated for language, using threshold Z>1.64, cluster threshold 10 voxels to identify suprathreshold voxels for the language contrast (Language>Control) in the left hemisphere language parcels and the mirror of these parcels in the right hemisphere. **(F)** As a point of comparison, we also calculated lateralization index for the social context contrast (Dialogue>Monologue) using the same approach and identical parcels (N=1 participant did not have suprathreshold voxels for this contrast).

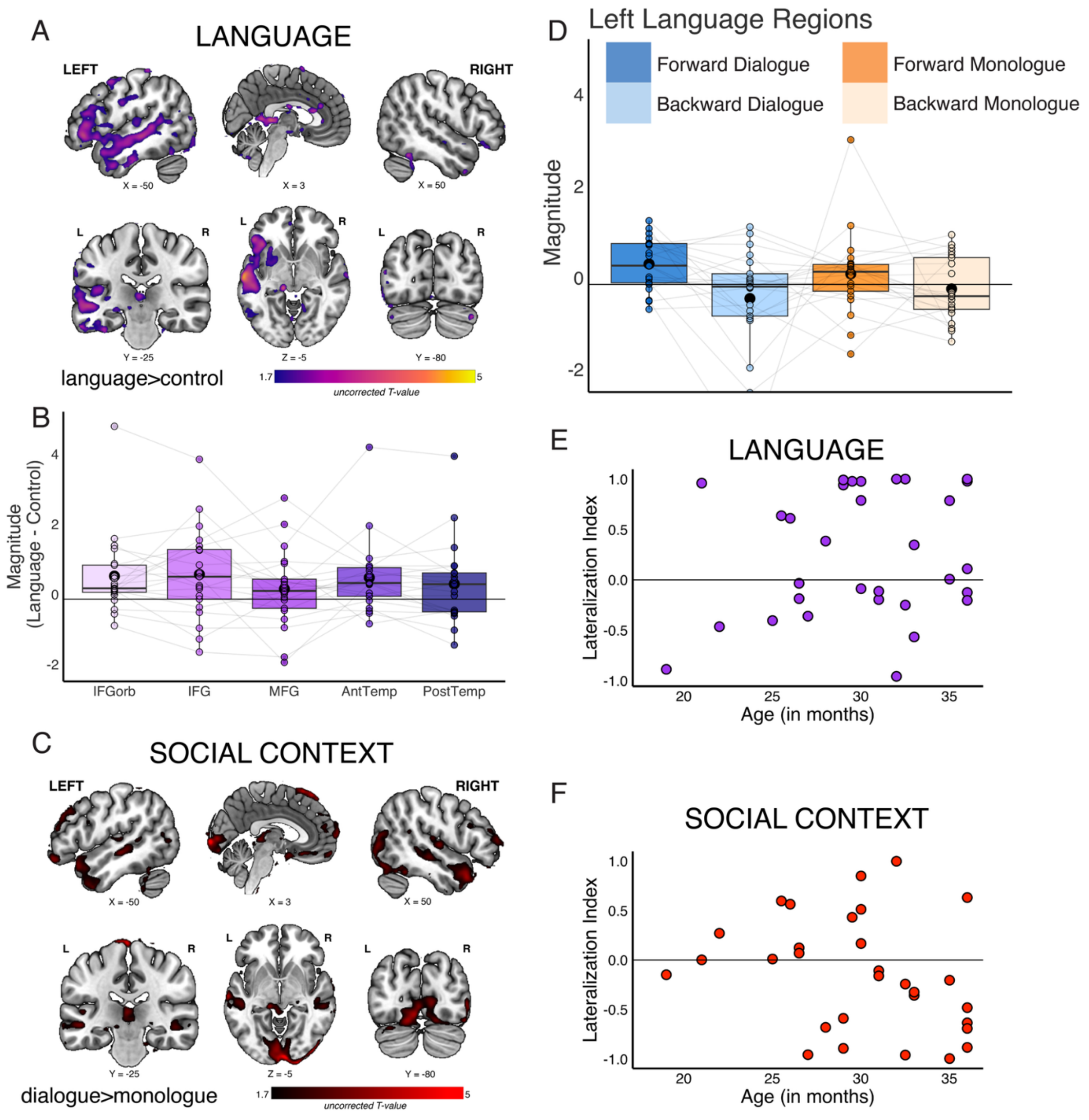

**Fig. S6.**

Results using infant-based motion cutoffs: we defined motion outliers as >3mm FD and excluded runs with >50% outliers. A group random effects analyses was conducted in the full sample of toddlers (N=31) with usable data at this threshold for **(A)** the language contrast and **(C)** the social context contrast. Toddler group responses are visualized at an uncorrected threshold T>1.70. **(D)** For participants with multiple usable functional runs (N=21), fROIs for the top 100 voxels were iteratively identified using the Language>Control contrast in each language parcel using a leave-one-run-out approach. Boxplots show response magnitude for held-out runs (independent data) for each condition averaged across all regions. **(B)** For each language region, we plot the difference in magnitude between language and control conditions in each individually-defined region (purple). Individual dots show each participant's response; light gray lines connect the same participant; large black dot shows mean. **(E)** For all 31 toddlers with usable data at this threshold, lateralization index was calculated for language, using threshold Z>1.64, cluster threshold 10 voxels to identify suprathreshold voxels for the language contrast

(Language>Control) in the left hemisphere language parcels and the mirror of these parcels in the right hemisphere. **(F)** As a point of comparison, we also calculated lateralization index for the social context contrast (Dialogue>Monologue) using the same approach and identical parcels (N=1 participant did not have suprathreshold voxels for this contrast).

(Language>Control) in the left hemisphere language parcels and the mirror of these parcels in the right hemisphere. **(F)** As a point of comparison, we also calculated lateralization index for the social context contrast (Dialogue>Monologue) using the same approach and identical parcels.

| | Included - Final Sample | Excluded - No Usable fMRI data |
|---|---|---|
| Sample size | N=29 | N=60 |
| Sex | 19 female / 10 male | 29 female / 31 male |
| Race/ethnicity | 19 white / 5 Asian / 1 other / 4 multi<br><br>4 Hispanic/Latino / 25 non-Hispanic/Latino | 23 white/ 10 Asian / 2 other / 17 multi, NA = 8<br><br>10 Hispanic/Latino / 43 non-Hispanic/Latino, NA= 7 |
| Gestational age (weeks) | mean(SD)=39.24(1.46) weeks, range=36-41 weeks | mean(SD)=39.13(1.60) weeks, range=34-41 weeks (NA=8) |
| Exposure to language(s) other than English | 14 English only / 15 multilingual | 17 English only / 31 multilingual (NA=12) |
| Average parental education | mean(SD)=18.46(1.85) years, range=14-22.5 years, NA=1 | mean(SD)=17.65(2.35) years, range=10-23 years, NA=12 |

**Table S1.**
Participant characteristics for the toddler sample included in analyses (N=29) and the remaining participants who were not included in analyses (N=60).

| Label (AAL Atlas) | Cluster Index | Voxels | MAX | MAX X (mm) | MAX Y (mm) | MAX Z (mm) | COG X (mm) | COG Y (mm) | COG Z (mm) |
|---|---|---|---|---|---|---|---|---|---|
| Temporal_Mid_L | 181 | 1311 | 3.63 | -62 | -20 | -4 | -54.9 | -18 | -5.7 |
| Temporal_Inf_L and Fusiform_L | 180 | 779 | 4.87 | -38 | -18 | -34 | -39.3 | -13.5 | -34.7 |
| | 179 | 358 | 4.5 | 0 | 22 | 16 | -1.78 | 27.4 | 13.1 |
| Temporal_Inf_R and Fusiform_R | 178 | 237 | 4.21 | 32 | 4 | -46 | 32.5 | 0.895 | -44.6 |
| Frontal_Inf_Orb_L | 177 | 230 | 3.1 | -40 | 26 | -4 | -44.2 | 28.9 | -3.99 |
| Cerebellum_Crus1_R | 176 | 156 | 3.23 | 54 | -50 | -38 | 49.7 | -47.8 | -28.7 |
| Precentral_R | 175 | 146 | 4.46 | 24 | -12 | 78 | 18.5 | -14.8 | 78.9 |
| Hippocampus_L | 174 | 139 | 3.11 | -18 | -12 | -14 | -21.5 | -14.9 | -12.5 |
| | 173 | 128 | 4.16 | -34 | -90 | -20 | -31.3 | -94.5 | -16.1 |
| Temporal_Mid_R and Temporal_Sup_R | 172 | 111 | 3.06 | 66 | -14 | -6 | 65.6 | -13.1 | -12.7 |
| Postcentral_L and SupraMarginal_L | 171 | 89 | 2.99 | -68 | -20 | 32 | -66.3 | -17.6 | 32.5 |
| Frontal_Sup_Medial_L | 170 | 80 | 2.62 | -2 | 38 | 34 | -1.42 | 34.3 | 33.4 |
| Supp_Motor_Area_R and Frontal_Sup_R | 169 | 69 | 3.07 | 6 | 8 | 74 | 8.95 | 5.8 | 74.9 |
| | 168 | 66 | 3.06 | 24 | -16 | -10 | 24.5 | -17.1 | -8.85 |
| | 167 | 56 | 2.98 | -42 | -86 | 30 | -50.5 | -77.2 | 31 |
| Cingulum_Ant_L | 166 | 53 | 2.22 | -2 | 44 | 4 | -3.08 | 45.7 | 5.02 |
| Precuneus_R | 165 | 52 | 2.73 | 14 | -48 | 78 | 9.38 | -44.4 | 77.8 |
| | 164 | 52 | 3.1 | 8 | -36 | -68 | 6.12 | -41.7 | -66.2 |
| Insula_L (tiny part in Putamen_L) | 163 | 51 | 2.16 | -34 | 10 | 4 | -36.1 | 10.3 | 4.54 |
| Temporal_Inf_L and Temporal_Mid_L | 162 | 49 | 2.91 | -62 | -60 | -6 | -63.3 | -54.6 | -6.26 |
| | 161 | 48 | 2.54 | 4 | -34 | 4 | 0.657 | -32.1 | 3.07 |
| Cerebelum_10_R | 160 | 45 | 2.66 | 26 | -32 | -38 | 26.9 | -30.9 | -35.8 |
| | 159 | 45 | 3.29 | 20 | -8 | -34 | 19.4 | -7.27 | -32.9 |
| | 158 | 38 | 2.22 | -34 | 12 | -18 | -33.7 | 12.3 | -17.4 |
| | 157 | 38 | 2.24 | 12 | 4 | -12 | 13.6 | 3.86 | -11.1 |
| | 156 | 37 | 3.38 | -68 | -34 | 38 | -67.9 | -33.9 | 37.6 |
| Precentral_L | 155 | 36 | 2.21 | -48 | 2 | 52 | -46.3 | 3.05 | 52.4 |
| Paracentral_Lobule_L | 154 | 34 | 3.29 | -16 | -24 | 80 | -14.5 | -25.5 | 79.8 |
| | 153 | 32 | 2.77 | -14 | 2 | 28 | -14.8 | 2.65 | 28.3 |
| Putamen_L | 152 | 31 | 2.23 | -18 | 10 | 8 | -17.4 | 10.4 | 6.91 |
| Temporal_Mid_R | 151 | 30 | 3.18 | 60 | 2 | -32 | 60.1 | 3.16 | -30.9 |
| Frontal_Sup_L | 150 | 30 | 1.99 | -18 | 38 | 34 | -16.7 | 40.3 | 34.5 |
| | 149 | 28 | 3.07 | -62 | -62 | 8 | -60.9 | -61.8 | 10.6 |
| Temporal_Pole_Sup_R | 148 | 28 | 2.88 | 46 | 8 | -12 | 46 | 8.25 | -12.5 |
| | 147 | 26 | 2.94 | -10 | -94 | -22 | -10.8 | -94.7 | -20.2 |
| Cerebelum_8_L | 146 | 26 | 2.65 | 36 | -38 | -44 | 31.5 | -37.9 | -46.3 |

| | | | | | | | | | |
|---|---|---|---|---|---|---|---|---|---|
| Postcentral_L and Temporal_Pole_Sup_L | 145 | 25 | 2.49 | -62 | -2 | 22 | -61 | -3.5 | 22.8 |
| Cerebelum_9_R | 144 | 25 | 2.11 | 8 | -54 | -38 | 7.84 | -53.7 | -38 |
| | 143 | 23 | 2.51 | -8 | 4 | -14 | -7.67 | 5.19 | -13 |
| Postcentral_L | 142 | 20 | 2.15 | -44 | -16 | 36 | -45.2 | -14.8 | 36.1 |
| Caudate_R | 141 | 20 | 2.11 | 16 | 14 | 4 | 13.8 | 12.9 | 3.52 |
| | 140 | 19 | 2.54 | -6 | -34 | -66 | -6.68 | -36.3 | -66.1 |
| | 139 | 19 | 3.31 | 40 | -80 | -34 | 39.1 | -79.4 | -33.1 |
| | 138 | 18 | 2.87 | -52 | -66 | 50 | -51.2 | -66.4 | 49.7 |
| | 137 | 16 | 2.77 | -56 | -72 | -10 | -54.9 | -71.6 | -11.8 |
| Temporal_Sup_R | 136 | 15 | 3.05 | 68 | -6 | 4 | 67.5 | -4.84 | 2.69 |
| Cerebelum_3_L | 135 | 14 | 2.09 | -12 | -34 | -24 | -12.1 | -33.6 | -24.2 |
| | 134 | 14 | 3.11 | 10 | -104 | 0 | 9.87 | -103 | -2.15 |
| Postcentral_L | 133 | 14 | 2.09 | -50 | -14 | 48 | -50.1 | -13.4 | 46.3 |
| Cerebelum_Crus1_L | 132 | 13 | 3 | -24 | -88 | -22 | -24 | -87.8 | -21.6 |
| Paracentral_Lobule_L | 131 | 13 | 3.14 | -6 | -28 | 80 | -4.52 | -29.5 | 80.5 |
| Frontal_Sup_L | 130 | 12 | 2.97 | -12 | 28 | 66 | -13.7 | 29.2 | 65.6 |
| Temporal_Pole_Mid_R | 129 | 12 | 2.02 | 60 | 4 | -16 | 58.2 | 4.99 | -14.5 |
| Temporal_Mid_R | 128 | 12 | 2.14 | 58 | -38 | 6 | 58.5 | -36.4 | 4.89 |
| Rolandic_Oper_R | 127 | 11 | 2.77 | 66 | 8 | 12 | 66 | 7.71 | 13.5 |
| Frontal_Inf_Tri_R | 126 | 11 | 2.48 | 64 | 22 | 12 | 64 | 22 | 9.51 |
| Temporal_Mid_R, some Temporal_Sup_R | 125 | 11 | 2.08 | 60 | -24 | -2 | 60.4 | -24.4 | -2.34 |
| Supp_Motor_Area_L | 124 | 11 | 2.78 | -14 | 14 | 72 | -12 | 14.3 | 72 |
| Parietal_Sup_R | 123 | 11 | 2.11 | 30 | -46 | -56 | 31.4 | -47.2 | -56.7 |
| Temporal_Mid_L and Temporal_Sup_L | 122 | 10 | 2.51 | -68 | -38 | -12 | -67.6 | -36.2 | -12 |
| Cingulum_Mid_L | 121 | 10 | 2.08 | 0 | 12 | 36 | 0.0136 | 13 | 36 |
| | 120 | 10 | 2.28 | -60 | -60 | 42 | -60.6 | -61.1 | 40.9 |
| Temporal_Sup_L | 119 | 10 | 2.04 | -42 | -26 | 14 | -43.3 | -26.2 | 13 |
| Frontal_Sup_L | 118 | 10 | 2.03 | -14 | -6 | 78 | -14.1 | -7.56 | 79 |
| | 117 | 10 | 2.32 | 4 | -78 | -46 | 3.86 | -76.1 | -44.7 |
| Cerebelum_Crus1_R | 116 | 10 | 1.91 | 20 | -88 | -20 | 19.6 | -87 | -21 |
| | 115 | 10 | 2.45 | 0 | -92 | -18 | -0.787 | -92.2 | -18.2 |

**Table S2.**

Significant clusters for the whole-brain group analyses (Language>Control), for all toddler participants (N=29). Uncorrected threshold, T>1.70; k>=10 voxels.

| Label (AAL Atlas) | Cluster Index | Voxels | MAX | MAX X (mm) | MAX Y (mm) | MAX Z (mm) | COG X (mm) | COG Y (mm) | COG Z (mm) |
|---|---|---|---|---|---|---|---|---|---|
| Lingual_R, Lingual_L, Calcarine_R, Calcarine_L | 183 | 6726 | 8.03 | 10 | -86 | 4 | 17.3 | -72.5 | 1.18 |
| Supp_Motor_Area_L, Frontal_Sup_Medial_L, Frontal_Sup_Medial_R, Supp_Motor_Area_R | 182 | 2444 | 6.29 | 12 | 20 | 70 | -1.95 | 4.38 | 70.4 |
| Frontal_Sup_Medial_L and Frontal_Sup_R | 181 | 1756 | 4.65 | -10 | 74 | 10 | -8.24 | 67.6 | 2.35 |
| Temporal_Mid_L | 180 | 1377 | 4.47 | -68 | -18 | -6 | -48.7 | 4.74 | -21.3 |
| Temporal_Pole_Sup_R and Temporal_Pole_Med_R | 179 | 1147 | 4.7 | 42 | 30 | -26 | 41 | 17.3 | -28.2 |
| Parietal_Inf_L | 178 | 1108 | 4.79 | -50 | -46 | 60 | -44.2 | -42.2 | 53.2 |
| Thalamus_L | 177 | 678 | 4.87 | 0 | -34 | 0 | -7.39 | -27.3 | 0.283 |
| Caudate_R | 176 | 656 | 3.39 | -6 | 6 | 6 | 12.2 | 11.9 | 2.87 |
| Cerelebum_8_L and Cerebelum_7b_L | 175 | 603 | 4.67 | -28 | -36 | -50 | -33.3 | -42.5 | -51.2 |
| Cerebelum_8_R | 174 | 462 | 3.61 | 38 | -58 | -60 | 29.2 | -52 | -56.4 |
| | 173 | 409 | 4.18 | 4 | -8 | -26 | 2.66 | -8.11 | -27.7 |
| Temporal_Mid_L | 172 | 377 | 2.97 | -44 | -72 | 4 | -49.8 | -55.8 | 10.7 |
| Precentral_L and Frontal_Inf_Oper_L, Frontal_Inf_Tri_L | 171 | 343 | 3.44 | -56 | 16 | 36 | -51.6 | 12.4 | 35.3 |
| | 170 | 215 | 3.57 | 64 | 16 | 0 | 60.9 | 24 | 6.55 |
| | 169 | 210 | 3.44 | 44 | -44 | 70 | 35.9 | -46.6 | 72.4 |
| Precuneus_R | 168 | 168 | 2.33 | 4 | -54 | 32 | 4.96 | -56.3 | 35 |
| Temporal_Sup_L and Heschl_L | 167 | 151 | 3.02 | -54 | -18 | 10 | -49.1 | -24.4 | 10.8 |
| Heschl_R and Temporal_Sup_R | 166 | 129 | 2.78 | 48 | -20 | 12 | 49.8 | -14.5 | 9.25 |
| Precentral_L | 165 | 115 | 2.56 | -26 | -10 | 48 | -30.2 | -7.72 | 49 |
| | 164 | 109 | 2.59 | 38 | -38 | 34 | 37.1 | -36.9 | 36.4 |
| Putamen_L and Caudate_L, a little Pallidum_L | 163 | 86 | 2.59 | -18 | 14 | 4 | -18.8 | 10.6 | 6.12 |
| | 162 | 86 | 3.03 | -14 | -8 | 26 | -14.7 | -3.93 | 26.9 |
| Supp_Motor_Area_L, Frontal_Sup_L, Supp_Motor_Area_R, Frontal_Sup_R, | 161 | 82 | 2.17 | 2 | 20 | 48 | 1.04 | 17.7 | 47.1 |
| Occipital_Sup_L | 160 | 79 | 3.08 | -12 | -92 | 32 | -13.6 | -89.2 | 30.5 |
| Cerebelum_Crus1_L, a little Temporal_Inf_R | 159 | 75 | 4.36 | -32 | -88 | -22 | -34.2 | -86.4 | -23.4 |
| Temporal_Pole_Sup_L, Temporal_Mid_L, | 158 | 64 | 3.9 | -38 | 6 | -20 | -38.6 | 5 | -21 |
| Hippocampus_R | 157 | 60 | 3.47 | 22 | -28 | -6 | 22.7 | -26.2 | -6.97 |

| | | | | | | | | | |
|---|---|---|---|---|---|---|---|---|---|
| Rectus_R, Rectus_L | 156 | 57 | 3.02 | -2 | 24 | -20 | 0.774 | 25.5 | -18.9 |
| | 155 | 54 | 3.41 | 26 | -40 | 14 | 26.9 | -41.1 | 12.4 |
| Occipital_Sup_L, Occipital_Mid_L | 154 | 54 | 2.95 | -10 | -102 | 10 | -12.7 | -102 | 9.83 |
| Occipital_Mid_L | 153 | 45 | 2.9 | -20 | -98 | 20 | -24.9 | -97.7 | 18.6 |
| | 152 | 44 | 2.52 | -28 | 18 | -8 | -26.4 | 19.2 | -8.18 |
| Frontal_Mid_R | 151 | 37 | 2.68 | 48 | 56 | 6 | 49.4 | 51.4 | 6.34 |
| Frontal_Inf_Tri_R | 150 | 37 | 2.35 | 34 | 16 | 26 | 34.1 | 16.2 | 25.7 |
| | 149 | 36 | 2.82 | 18 | -26 | 24 | 18.4 | -29.6 | 24 |
| Temporal_Inf_R | 148 | 35 | 2.26 | 50 | -22 | -26 | 52.9 | -20.3 | -28.7 |
| Frontal_Mid_L | 147 | 31 | 2.87 | -46 | 16 | 52 | -44 | 15.7 | 53.4 |
| ParaHippocampal_R | 146 | 30 | 2.57 | 18 | 2 | -20 | 18.9 | 2.61 | -20.4 |
| Precentral_L | 145 | 28 | 2.32 | -46 | 6 | 18 | -46.3 | 5.65 | 18.9 |
| Pallidum_R | 144 | 27 | 2.15 | 20 | -2 | 8 | 18.5 | -0.456 | 6.02 |
| | 143 | 26 | 2.39 | -14 | -18 | 28 | -15.6 | -21.1 | 27.1 |
| | 142 | 25 | 2.82 | 2 | 32 | 8 | 1.45 | 32.6 | 7.43 |
| | 141 | 22 | 2.33 | -24 | -44 | 14 | -25.2 | -45.5 | 12.1 |
| Lingual_R, Calcarine_R, Lingual_L, Calcarine_L | 140 | 21 | 2.35 | 22 | -56 | -2 | 21.4 | -57.5 | -2.9 |
| Precentral_R | 139 | 20 | 1.95 | 36 | 6 | 32 | 38.6 | 3.91 | 32.6 |
| | 138 | 20 | 2.34 | 52 | 38 | -18 | 50.6 | 42.3 | -17.9 |
| Frontal_Mid_L | 137 | 20 | 2.86 | -40 | 34 | 46 | -44 | 30.6 | 44.8 |
| Occipital_Sup_R | 136 | 19 | 2.36 | 12 | -86 | 40 | 15.4 | -87.9 | 36.1 |
| Cingulum_Ant_R | 135 | 19 | 2.39 | 4 | 54 | 16 | 3.16 | 54.2 | 15.4 |
| Parietal_Inf_R | 134 | 18 | 2.89 | 58 | -42 | 56 | 56.6 | -42.9 | 55.1 |
| Frontal_Mid_L | 133 | 17 | 2.5 | -48 | 42 | 18 | -49.5 | 41.3 | 18.3 |
| | 132 | 15 | 2.32 | -48 | -88 | 4 | -47.7 | -87.2 | 4.31 |
| Frontal_Sup_Medial_R, Supp_Motor_Area_R, Supp_Motor_Area_L, Frontal_Sup_L | 131 | 15 | 2.16 | 4 | 50 | 34 | 4 | 49.6 | 34.8 |
| Cuneus_L | 130 | 14 | 2.47 | 4 | -82 | 38 | 2.01 | -84 | 37.5 |
| Postcentral_R | 129 | 13 | 2.28 | 18 | -36 | 82 | 18.5 | -36.1 | 81.4 |
| Fusiform_L | 128 | 13 | 2.32 | -36 | -40 | -24 | -36.3 | -39.9 | -23.6 |
| Cerebelum_Crus2_R | 127 | 13 | 2.51 | 42 | -84 | -38 | 42.7 | -82.2 | -38 |
| Cerelebum_9_R | 126 | 12 | 2.03 | 2 | -56 | -54 | 1.77 | -55.2 | -54.5 |
| | 125 | 12 | 2.18 | 36 | -18 | 36 | 36 | -19 | 35.6 |
| Lingual_L, Calcarine_L, Lingual_R, Calcarine_R | 124 | 12 | 2.17 | -18 | -52 | -2 | -19.3 | -53.5 | -1.2 |

| | | | | | | | | | |
|---|---|---|---|---|---|---|---|---|---|
| Frontal_Inf_Tri_L | 123 | 11 | 2.17 | -50 | 22 | 14 | -50.6 | 21.8 | 14.5 |
| Fusiform_R | 122 | 11 | 2.33 | 40 | -20 | -32 | 42.1 | -22.4 | -30.7 |
| Temporal_Sup_R | 121 | 11 | 2.15 | 48 | -6 | -6 | 48.7 | -5.82 | -6 |
| Postcentral_L | 120 | 10 | 2.03 | -56 | -18 | 32 | -56 | -18 | 33 |
| | 119 | 10 | 2.32 | 50 | -6 | -42 | 49.2 | -5.63 | -43.5 |
| Cerebelum_Crus2_R | 118 | 10 | 2.27 | 32 | -84 | -42 | 34 | -82.4 | -42.8 |

**Table S3.**

Significant clusters for the whole-brain group analyses (Dialogue>Monologue), for all toddler participants (N=29). Uncorrected threshold, T>1.70; k>=10 voxels.

| Network | Forward v. Backward | Dialogue v. Monologue | Temporal v. Frontal | Interactions | Age |
|---|---|---|---|---|---|
| **Toddlers: Left Language** | Est=0.175<br>S.E.=0.050<br>t-value=3.492<br>**p<0.001 *** | Est=0.094<br>S.E.=0.050<br>t-value=1.869<br>p=0.062 | Est=0.162<br>S.E.=0.060<br>t-value=2.702<br>**p=0.047 *** | None significant | Est.=0.005<br>S.E.=0.024<br>t-value=0.200<br>p=0.844 |
| **Toddlers: Right Language** | Est=0.065<br>S.E.=0.050<br>t-value=1.296<br>p=0.196 | Est=0.049<br>S.E.=0.050<br>t-value=0.980<br>p=0.328 | Est=0.160<br>S.E.=0.050<br>t-value=3.195<br>**p=0.0015 *** | None significant | Est.=0.025<br>S.E.=0.026<br>t-value=0.982<br>p= 0.340 |
| **Adults: Left Language** | Est.= 1.266<br>S.E.= 0.047<br>t-value= 27.162<br>**p<0.001 *** | Est.= 0.092<br>S.E.= 0.047<br>t-value= 1.983<br>**p= 0.0481 *** | Est.= 0.687<br>S.E.= 0.161<br>t-value= 4.255<br>**p= 0.009 *** | f_or_b*t_or_f:<br>Est.= 0.375<br>S.E.= 0.047<br>t-value= 8.045<br>**p<0.001 ***<br><br>*Others not significant* | *Not included in model* |
| **Adults: Right Language** | Est.= 0.873<br>S.E.= 0.056<br>t-value= 15.705<br>**p<0.001 *** | Est.= 0.203<br>S.E.= 0.056<br>t-value= 3.652<br>**p<0.001 *** | Est.= 1.000<br>S.E.= 0.213<br>t-value= 4.692<br>**p= 0.006 *** | f_or_b*t_or_f:<br>Est.= 0.436<br>S.E.= 0.056<br>t-value= 7.832<br>**p<0.001 ***<br><br>*Others not significant* | *Not included in model* |

**Table S4.**

Model results across left-hemisphere language network (5 fROIs) and right-hemisphere homotopes of language network (5 fROIs) in toddlers (N=17) and adults (N=20). FROIs defined as top 100 voxels. Model: lmer(avg_beta~f_or_b *d_or_m*t_or_f+age+(1|participantID)+(1|ROI), REML = FALSE). Note that age was not included in the adult models.
* indicates significance level p<0.05

| ROI | Forward v. Backward | Dialogue v. Monologue | Interaction (f_or_b*d_or_m) | Age |
|---|---|---|---|---|
| Left IFGorb | Est.=0.288<br>S.E.=0.104<br>t-value=2.774<br>**p=0.0077 *** | Est.=0.110<br>S.E.=0.104<br>t-value=1.062<br>p=0.293 | Est.=0.065<br>S.E.=0.104<br>t-value=0.623<br>p=0.536 | Est.=0.024<br>S.E.=0.028<br>t-value=0.845<br>p=0.410 |
| Left IFG | Est.=0.242<br>S.E.=0.110<br>t-value=2.200<br>p=0.0324 | Est.=0.0068<br>S.E.=0.110<br>t-value=0.062<br>p=0.9509 | Est.=0.0004<br>S.E.=0.110<br>t-value=0.004<br>p=0.997 | Est.=-0.044<br>S.E.=0.030<br>t-value=-1.477<br>p=0.158 |
| Left MFG | Est.=0.047<br>S.E.=0.094<br>t-value=0.497<br>p=0.621 | Est.=0.132<br>S.E.=0.094<br>t-value=1.414<br>p=0.163 | Est.=0.153<br>S.E.=0.094<br>t-value=1.631<br>p=0.109 | Est.=0.018<br>S.E.=0.033<br>t-value=0.561<br>p=0.582 |
| Left AntTemp | Est.=0.204<br>S.E.=0.064<br>t-value=3.171<br>**p=0.0026 *** | Est.=0.093<br>S.E.=0.064<br>t-value=1.441<br>p=0.156 | Est.=-0.062<br>S.E.=0.064<br>t-value=-0.972<br>p=0.336 | Est.=0.040<br>S.E.=0.043<br>t-value=0.937<br>p=0.362 |
| Left PostTemp | Est.=0.112<br>S.E.=0.097<br>t-value=1.146<br>p=0.257 | Est.=0.116<br>S.E.=0.097<br>t-value=1.189<br>p=0.240 | Est.=0.095<br>S.E.=0.097<br>t-value=0.970<br>p=0.337 | Est.=-0.015<br>S.E.=0.052<br>t-value=-0.291<br>p=0.774 |

**Table S5.**

Toddler model results (N=17) for each individual left-hemisphere language parcel, defined as top 100 voxels. Model: lmer(avg_beta~f_or_b *d_or_m+age+(1|participantID), REML = FALSE)
* indicates significance level p<0.05, Bonferroni corrected for 5 ROIs (p<0.01).

| ROI | Forward v. Backward | Dialogue v. Monologue | Interaction (f_or_b*d_or_m) |
|---|---|---|---|
| Left IFGorb | Est.=0.787<br>S.E.= 0.072<br>t-value= 10.929<br>**p<0.001 *** | Est.= -0.007<br>S.E.= 0.072<br>t-value= -0.097<br>p= 0.923 | Est.= 0.031<br>S.E.= 0.072<br>t-value= 0.432<br>p= 0.667 |
| Left IFG | Est.= 0.952<br>S.E.= 0.066<br>t-value= 14.390<br>**p<0.001 *** | Est.= 0.026<br>S.E.= 0.066<br>t-value= 0.396<br>p= 0.693 | Est.= 0.011<br>S.E.= 0.066<br>t-value= 0.162<br>p= 0.872 |
| Left MFG | Est.= 0.934<br>S.E.= 0.073<br>t-value=12.856<br>**p<0.001 *** | Est.= 0.061<br>S.E.= 0.073<br>t-value= 0.833<br>p= 0.408 | Est.= 0.027<br>S.E.= 0.073<br>t-value= 0.378<br>p= 0.707 |
| Left AntTemp | Est.= 1.392<br>S.E.= 0.059<br>t-value= 23.404<br>**p<0.001 *** | Est.= 0.145<br>S.E.= 0.059<br>t-value= 2.443<br>p= 0.0175 | Est.= 0.064<br>S.E.= 0.059<br>t-value= 1.071<br>p= 0.288 |
| Left PostTemp | Est.= 1.891<br>S.E.= 0.070<br>t-value=27.153<br>**p<0.001 *** | Est.= 0.171<br>S.E.= 0.070<br>t-value= 2.460<br>p= 0.0168 | Est.= 0.038<br>S.E.= 0.070<br>t-value= 0.544<br>p= 0.588 |

**Table S6.**

Adult model results (N=20) for each individual left-hemisphere language parcel, defined as top 100 voxels. Model: lmer(avg_beta~f_or_b *d_or_m+ (1|participantID), REML = FALSE)
* indicates significance level p<0.05, Bonferroni corrected for 5 ROIs (p<0.01).

| ROI | Forward v. Backward | Dialogue v. Monologue | Interaction (f_or_b*d_or_m) | Age |
|---|---|---|---|---|
| Right IFGorb | Est.=0.116<br>S.E.=0.118<br>t-value=0.984<br>p=0.330 | Est.=0.095<br>S.E.=0.118<br>t-value=0.802<br>p=0.426 | Est.=0.013<br>S.E.=0.118<br>t-value=0.109<br>p=0.914 | Est.= 0.002<br>S.E.= 0.037<br>t-value= 0.042<br>p= 0.967 |
| Right IFG | Est.=-0.063<br>S.E.=0.089<br>t-value=-0.706<br>p=0.483 | Est.=0.069<br>S.E.=0.089<br>t-value=0.775<br>p=0.442 | Est.=-0.098<br>S.E.=0.089<br>t-value=-1.100<br>p=0.276 | Est.= -0.008<br>S.E.= 0.055<br>t-value= -0.152<br>p= 0.881 |
| Right MFG | Est.=-0.049<br>S.E.=0.088<br>t-value=-0.560<br>p=0.578 | Est.=0.071<br>S.E.=0.088<br>t-value=0.807<br>p=0.423 | Est.=0.164<br>S.E.=0.088<br>t-value=1.859<br>p=0.069 | Est.= -0.009<br>S.E.= 0.025<br>t-value= -0.369<br>p= 0.717 |
| Right AntTemp | Est.=0.193<br>S.E.=0.090<br>t-value=2.140<br>p=0.0371 | Est.=0.072<br>S.E.=0.090<br>t-value=0.794<br>p=0.4311 | Est.=0.008<br>S.E.=0.090<br>t-value=0.084<br>p=0.9333 | Est.= 0.071<br>S.E.= 0.036<br>t-value= 1.978<br>p= 0.064 |
| Right PostTemp | Est.=0.064<br>S.E.=0.090<br>t-value=0.714<br>p=0.478 | Est.=-0.032<br>S.E.=0.090<br>t-value=-0.350<br>p=0.728 | Est.=-0.093<br>S.E.=0.090<br>t-value=-1.036<br>p=0.305 | Est.= 0.071<br>S.E.= 0.036<br>t-value= 1.949<br>p= 0.068 |

**Table S7.**

In addition to measuring responses in canonical left-hemisphere language parcels, we extracted responses in individually-defined fROIs (top 100 voxels) for the Language>Control contrast in mirror parcels in the right hemisphere (right-hemisphere homotopes). Toddler model results (N=17) for each individual right-hemisphere language parcel. * indicates significance level p<0.05, Bonferroni corrected for 5 ROIs (p<0.01).

| Threshold | # voxels - left | # voxels - right | LI | One-sample t-test: LI>0 |
|---|---|---|---|---|
| z = 2.58 | M(SD)= 97.03(182.01); range=0-739 | M(SD)= 46.45(105.54); range=0-532 | M(SD)= 0.264(0.77); range=-1-1, NA=10 | t= 1.49; p=0.077 |
| z = 2.33 | M(SD)= 172.4(277.38); range=0-1155 | M(SD)= 82(170.37); range=0-823 | M(SD)= 0.29(0.73); range=-1-1; NA=6 | t=1.92; **p=0.033 *** |
| z = 1.96 | M(SD)= 358.6(469.23); range=0-1821 | M(SD)= 181.9(327.76); range=0-1461 | M(SD)= 0.288(0.68); range=-1-1 | t= 2.28; **p=0.015 *** |
| z = 1.64 | M(SD)= 599(699.23); range=0-2399 | M(SD)= 333.4(515.43); range=0-2127 | M(SD)= 0.258(0.61); range=-1-1 | t=2.28; **p=0.015 *** |
| z = 1.28 | M(SD)= 1010(936.66); range=1-3093 | M(SD)= 614.8(778.29); range=0-3125 | M(SD)= 0.234(0.56); range=-0.98-1 | t=2.24; **p=0.016 *** |
| z = 0.84 | M(SD)= 1739(1279.50); range=26-4323 | M(SD)= 1175(1112.92); range=16-4359 | M(SD)= 0.182(0.50); range=-0.89-0.96 | t=1.94; **p=0.031 *** |

**Table S8.**
Toddler (N=29) lateralization index (LI) calculated at different thresholds for identifying significant language voxels (Forward>Backward contrast; cluster threshold k>=10 voxels). Number of voxels in the left and right search spaces (5 parcels) are reported.

| ROI | Forward v. Backward | Dialogue v. Monologue | Interaction (f_or_b*d_or_m) | Age |
|---|---|---|---|---|
| Left IFGorb | Est.=0.317<br>S.E.=0.114<br>t-value=2.771<br>**p=0.008 *** | Est.=0.074<br>S.E.=0.114<br>t-value=0.644<br>p=0.523 | Est.=0.087<br>S.E.=0.114<br>t-value=0.759<br>p=0.452 | Est.=0.036<br>S.E.=0.032<br>t-value=1.127<br>p=0.275 |
| Left IFG | Est.=0.253<br>S.E.=0.113<br>t-value=2.236<br>p=0.0297 | Est.=0.004<br>S.E.=0.113<br>t-value=0.036<br>p=0.972 | Est.=-0.006<br>S.E.=0.113<br>t-value=-0.056<br>p=0.956 | Est.=-0.045<br>S.E.=0.031<br>t-value=-1.416<br>p=0.175 |
| Left MFG | Est.=0.012<br>S.E.=0.098<br>t-value=0.117<br>p=0.907 | Est.=0.142<br>S.E.=0.098<br>t-value=1.449<br>p=0.154 | Est.=0.144<br>S.E.=0.098<br>t-value=1.466<br>p=0.149 | Est.=0.035<br>S.E.=0.034<br>t-value=1.023<br>p=0.321 |
| Left AntTemp | Est.=0.193<br>S.E.=0.060<br>t-value=3.224<br>**p=0.002 *** | Est.=0.102<br>S.E.=0.060<br>t-value=1.700<br>p=0.095 | Est.=-0.047<br>S.E.=0.060<br>t-value=-0.778<br>p=0.440 | Est.=0.040<br>S.E.=0.040<br>t-value=0.992<br>p=0.335 |
| Left PostTemp | Est.=0.118<br>S.E.=0.088<br>t-value=1.341<br>p=0.186 | Est.=0.109<br>S.E.=0.088<br>t-value=1.242<br>p=0.220 | Est.=0.069<br>S.E.=0.088<br>t-value=0.787<br>p=0.435 | Est.=-0.009<br>S.E.=0.044<br>t-value=-0.198<br>p=0.845 |

**Table S9.**

Toddler model results (N=17) for each individual left-hemisphere language parcel, defined as top 10% of voxels within each search space. * indicates significance level p<0.05, Bonferroni corrected for 5 ROIs (p<0.01).
